# Lactate epigenetically shapes tumor cell states in H3K27M diffuse midline gliomas

**DOI:** 10.64898/2026.10.02.756321

**Authors:** Joanna Lum, Nicholas Nuechterlein, Shelby Sliger, Sanjana Eyunni, Benita Tamrazi, Stefan Blüml, Wentao Tian, Akash Deogharkar, Stefan R. Sweha, Mateus Mota, Jill Bayliss, Erik Peterson, Sunjong Ji, Molly Wittwer, Ksheetesha Bhat, Bryan T. Mott, Erin T. Hamanishi, Fjodor Merkuri, Ana Azambuja, Helena Conceicao, Fengyun Su, Xuhong Cao, Rishikesh Pandey, Omkar B. Ljare, Abhijit Parolia, Viveka Nand Yadav, Daniel R Wahl, Carl Koschmann, Marcos Simoes-Costa, Kumar Pichumani, Shaomeng Wang, Arul M Chinnaiyan, Sriram Venneti

**Affiliations:** Laboratory of Brain Tumor Metabolism and Epigenetics, Department of Pathology, University of Michigan, Ann Arbor, MI 48109, USA; Chad Carr Pediatric Brain Tumor Center, Department of Pediatrics, University of Michigan, Ann Arbor, MI, USA; Rogel Cancer Center, University of Michigan Medical School, Ann Arbor, MI, USA; Department of Pathology, University of Michigan, Ann Arbor, MI 48109; Michigan Center for Translational Pathology, University of Michigan, Ann Arbor, MI 48109; Department of Radiology, Children’s Hospital Los Angeles, Keck School of Medicine, University of Southern California, Los Angeles, CA 90027, USA; Memorial Sloan Kettering Cancer Center, Weill Cornell Graduate School of Medical Sciences, New York, NY 10065, USA; Department of Radiation Oncology, University of Michigan, Ann Arbor, MI, USA; Department of Pediatrics, Division of Pediatric Hematology-Oncology, University of Michigan School of Medicine, Ann Arbor, MI, USA; Department of Neurological Surgery, Wake Forest Baptist Medical Center, Winston-Salem, NC 27157, USA; Division of Hematology/Oncology, Department of Pediatrics, The Hospital for Sick Children and University of Toronto, Toronto, ON, Canada; Department of Pathology, Boston Children’s Hospital, Boston, MA, USA; Department of Systems Biology, Harvard Medical School, Boston, MA, USA; Howard Hughes Medical Institute, University of Michigan, Ann Arbor, MI, USA; Kenneth R. Peak Brain and Pituitary Tumor Treatment Center, Department of Neurosurgery, Houston Methodist Hospital, Houston, Texas 77030, USA; Houston Methodist Research Institute, Houston, Texas 77030, USA; Department of Urology, University of Michigan, Ann Arbor Arbor, MI, USA; Center for Molecular Oncology and Cellular Therapeutics (MOCT), Children’s Mercy Research Institute (CMRI), and Department of Pediatrics, University of Missouri– Kansas City School of Medicine, Kansas City, MO 64108, USA; Weill Cornell Medical College, New York, New York 10065, USA; Department of Internal Medicine, Division of Hematology/Oncology, University of Michigan, Ann Arbor, Michigan 48109, United States; Department of Pharmacology, Medical School, University of Michigan, Ann Arbor, Michigan 48109, United States; Department of Medicinal Chemistry, College of Pharmacy, University of Michigan, Ann Arbor, Michigan 48109, United States; Department of Computational Medicine and Bioinformatics, University of Michigan, Ann Arbor, MI, USA

## Abstract

H3K27M diffuse midline gliomas (DMGs) are lethal and lack treatment options. These tumors exhibit tumor heterogeneity with aggressive oligodendrocyte precursor (OPC)- and mesenchymal (MES)-like cell states. Because all tumor cells bear H3K27M, tumor heterogeneity has been attributed to microenvironmental cues that remain obscure. H3K27M DMGs exhibit increased glycolysis and the Warburg effect to produce lactate. Lactate can directly modify histone lysines including H3K27Lac, and we hypothesized that lactate epigenetically regulates tumor cell states in H3K27M DMGs. We undertook a comprehensive and integrated analysis of H3K27M DMG cell lines, animal models, tumor tissues, and magnetic resonance spectroscopy in patients to decipher the epigenetic roles of lactate. Here, we report that H3K27M cells uniquely exhibit elevation of the activating mark H3K27Lac. We observed significant heterogeneity in lactate levels within tumors, and the addition of exogenous lactate resulted in marked increases in H3K27Lac and the expression of critical OPC- and MES- like gene signature genes, which correlates with poorer patient outcomes. Our therapeutic approach leverages the inhibition of CBP/p300, which deposits both H3K27Lac and H3K27ac. Targeting CBP/p300 effectively reduces both H3K27Lac and H3K27ac levels, suppresses key OPC- and MES-like signatures, and shows efficacy in preclinical models. Collectively, our findings identify lactate as a key player in the epigenetic regulation of tumor cell state heterogeneity and suggest that targeting this pathway offers promise for addressing the heterogeneous complexities of H3K27M DMGs.

## Introduction

H3K27M Diffuse midline gliomas (DMGs) are aggressive grade 4 tumors arising from midline structures including the thalamus, midbrain, or pons.^1^ Due to their location and diffusely infiltrative features, these tumors are poorly circumscribed and generally not operable. DMGs primarily affect children ages 5-9 years old, lack curative therapy, and have a dismal prognosis of 9-11 months post-diagnosis.^2,3^ These tumors are defined by the H3K27M (including H3.1 and H3.3) histone mutation.^4–6^ H3K27M are missense mutations that substitute lysine (K) 27 with methionine (M), leading to global loss of H3K27me3.^7–10^ Activating H3K27ac, which opposes H3K27me3, is globally increased and enriched at lineage related genes, but shows widespread deposition at intergenic regions including endogenous retroviral elements.^11–15^ This is accompanied by extensive epigenetic reprogramming including global DNA hypomethylation, aberrant bivalent marks, H3K36 methylation, and altered chromatin.^7,16–22^

Single cell multi-omics and spatial transcriptomics of H3K27M DMGs tissues revealed complex heterogeneity in cellular composition across age and location.^23–29^ Tumor cells show oligodendrocyte (OC)-like, astrocyte (AC)-like, oligodendrocyte precursor cell (OPC)-like, and mesenchymal (MES)-like cells, with OPC-like and MES-like cells displaying high malignant potential.^23–29^ Because all tumor cells bear H3K27M mutations, tumor heterogeneity cannot be entirely attributed to the mutation. This has led to the long-standing hypothesis that microenvironmental cues play a key role in shaping tumor heterogeneity. However, specific environmental cues that drive this phenomenon remain poorly understood. Addressing this major gap in our knowledge is critical to understand tumor heterogeneity in therapeutic response and development of treatment resistance.

Metabolic reprogramming is a hallmark of cancer.^30^ H3K27M mutant gliomas demonstrate aberrant metabolism.^31–36^ In H3K27M cells, metabolic rewiring is closely linked with epigenetic dysregulation and cell fate.^32,36,37^ Recently, the metabolite, lactate, was found to be able to decorate lysine (K) residues on histones, a process termed histone lactylation.^38,39^ In the broader context, this discovery directly connects the Warburg effect and lactate production with transcriptional regulation. Because integrated metabolic and epigenetic pathways govern cell fate, we hypothesized that the metabolite lactate epigenetically modulates tumor heterogeneity in H3K27M DMGs. Based on this premise, we assessed if histone lactylation is a functionally important process in DMG biology.

## Results

### H3K27Lac is higher in H3K27M versus H3WT cells

Because H3K27M cells exhibit the Warburg effect and show high lactate levels,^32,40–43^ we assessed for differences in histone lactylation between isogenic cells expressing H3.3K27M or H3WT in immortalized mouse neuronal stem cells (mNSCs).^32^ H3.3K27M versus H3WT mNSC showed globally reduced H3K27me3, accompanied by increased H3K27ac (**Fig 1A**), as shown by others.^11,12,14,15,21^ We profiled ten different histone lactylation modifications at lysine residues in H2A, H2B, H3, and H4. H3K27Lac was higher in H3.3K27M versus H3WT mNSCs (**Fig 1A**). All other marks including H3K9Lac, H3K14Lac, H3K18Lac, and other H2AK-, H2BK-, H4K- lactyl marks were not different between H3.3K27M versus H3WT mNSCs (**Fig 1A, Fig S1A-C**). We determined if H3K27Lac levels were different in another histone mutant glioma model: H3G34-mutant diffuse high-grade glioma. We did not see any consistent differences in H3K27Lac or other histone K-lactylation marks between H3.3G34R versus H3WT expressing mNSCs (**Fig S1D-E**).

**Figure 1.**
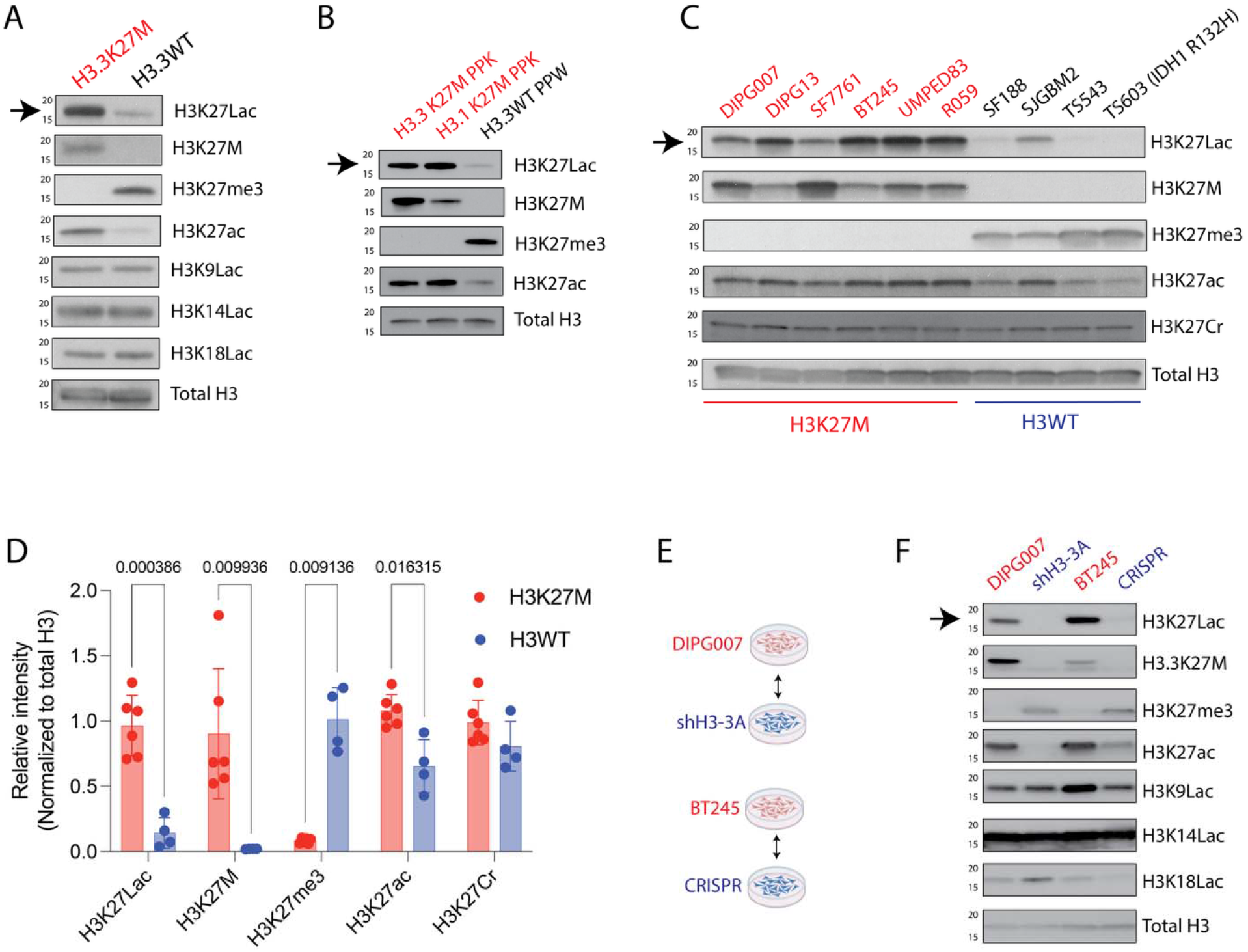
H3K27Lac is higher in H3K27M versus H3WT cells. - **(A)** Representative Western blots of H3K27Lac (arrow), H3K27M, H3K27me3, H3K27ac, H3K9Lac, H3K14Lac, H3K18Lac, and total H3 in H3.3K27M (red) and H3.3WT (blue) mouse neuronal stem cells (mNSCs).
- **(B)** Representative Western blots of H3K27Lac (arrow), H3K27M, H3K27me3, H3K27ac, and total H3 in H3.3K27M (red) PPK, H3.1K27M (red) PPK, and H3.3WT (blue) PPW cells. PPK/PPW *in utero* electroporation derived PdgfrA-overexpressing and shp53, with H3.3K27M or H3.1K27M or H3WT conditions.
- **(C)** Representative Western blots of H3K27Lac (arrow), H3K27M, H3K27me3, H3K27ac, H3K27Cr, and total H3 in H3.3K27M (DIPG007, DIPG13, SF7761, BT245, UMPED83, R059: red) and H3WT (SF188, SJGBM2, TS543, and TS603: blue) patient-derived cell lines.
- **(D)** Relative Western blots intensity (normalized to total H3) of H3K27Lac, H3K27M, H3K27me3, H3K27ac, and H3K27Cr from panel of H3K27M and H3WT cells lines shown in main figure 1C. Data shown as mean ± SD and analyzed by using two-sided, unpaired, two tailed **t** test with 95% C.I. P values indicated in figure.
- **(E)** Diagram showing the generation of isogenic shH3-3A (shRNA-mediated stable knockdown of *H3-3A*) and CRISPR (CRISPR-mediated correction of K-M) lines from their corresponding DIPG007 and BT245 parental cells.
- **(F)** Representative Western blots of H3K27Lac, H3.3K27M, H3K27me3, H3K27ac, H3K9Lac, H3K14Lac, H3K18Lac, and total H3 in DIPG007 (red) versus DIPG007-shH3-3A (blue), and BT245 (red) versus BT245-CRISPR (blue) cell lines. Figures in panel E were generated using biorender.

**Figure S1.**
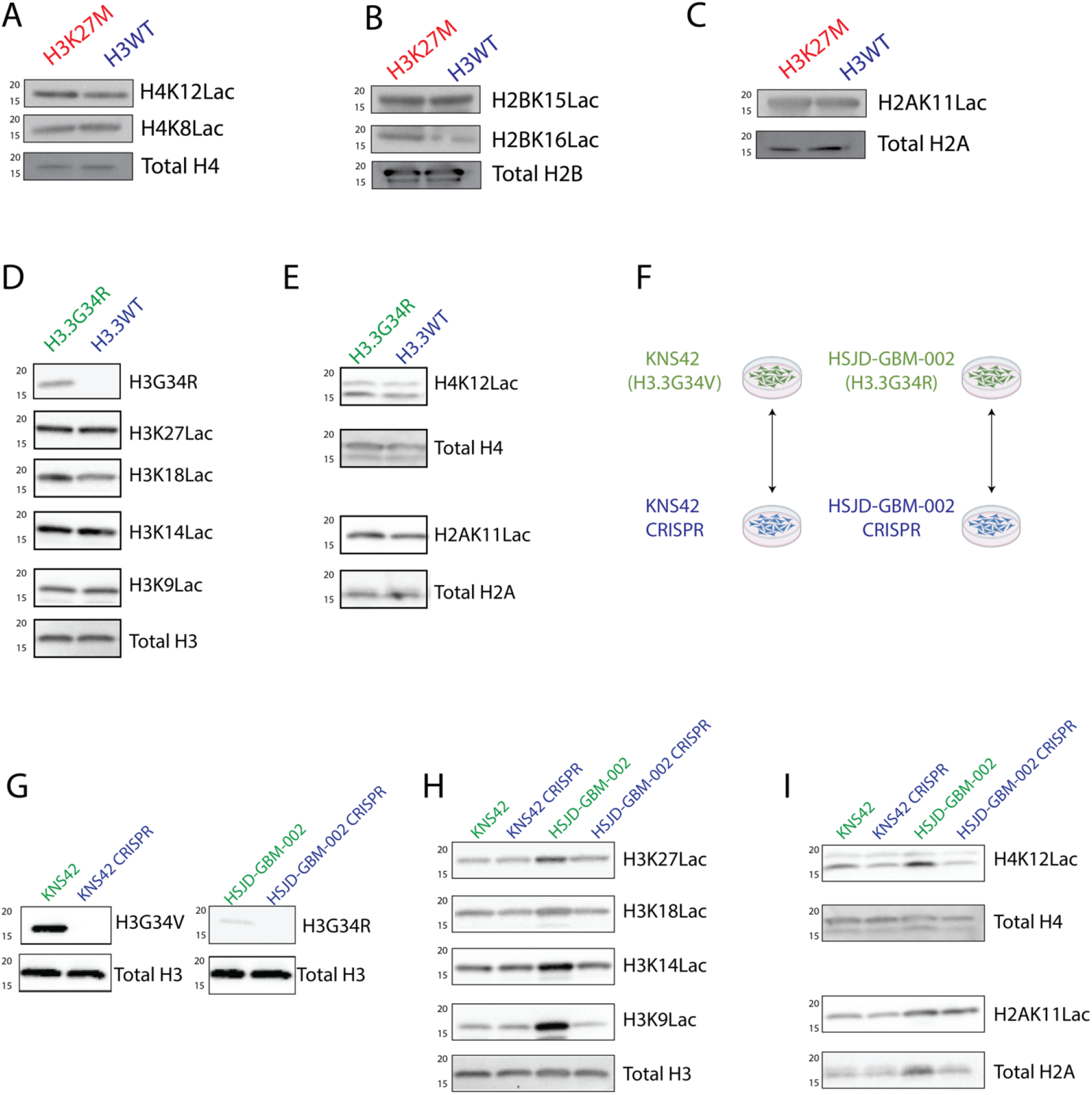
H3K27Lac is higher in H3K27M versus H3WT cells (related to figure 1) - **(A)** Representative Western blots of H4K12Lac, H4K8Lac, and total H4 in H3.3K27M and H3.3WT mNSCs.
- **(B)** Representative Western blots of H2BK15Lac, H2BK16Lac and total H2B in H3.3K27M and H3.3WT mNSCs.
- **(C)** Representative Western blots of H2AK11Lac and total H2A in H3.3K27M and H3.3WT mNSCs.
- **(D)** Representative western blots of H3K27Lac, H3K18Lac, H3K14Lac, H3K9Lac, and total H3 in H3.3G34R and H3.3WT mNSCs.
- **(E)** Representative Western blots of H4K12Lac, total H4, H2AK11Lac, and total H2A in H3.3G34R and H3.3WT mNSCs.
- **(F)** Diagram showing isogenic KNS42 H3.3G34V mutant cells with or without CRISPR V-G correction; and HSJD-GBM-002 H3.3G34R mutant cells with or without CRISPR R-G correction.
- **(G)** Representative Western blots of mutant specific H3G34R, H3G34V, and total H3 in isogenic KNS42 versus CRISPR (left), and HSJD-GBM-002 versus CRISPR (right) cell lines.
- **(H)** Representative Western blots of H3K27Lac, H3K18Lac, H3K14Lac, H3K9Lac, and total H3 in isogenic KNS42 versus CRISPR, and HSJD-GBM-002 versus CRISPR cell lines.
- **(I)** Representative Western blots of H4K12Lac, total H4, H2AK11Lac, and total H2A in in isogenic KNS42 versus CRISPR, and HSJD-GBM-002 versus CRISPR cell lines. Figures in panel F were generated using biorender.

Since H3K27Lac was the only histone lactylation mark elevated in H3K27M versus H3WT mNSC, we validated our finding in a wider panel of cells. We used H3.3K27M, H3.1K27M, versus H3WT cells derived from *in utero* electroporation (IUE) models (PPK/PPW: PdgfrA-overexpressed and shp53 with H3.3K27M or H3.1K27M or H3WT conditions, **Fig 1B**). We also assessed a panel of H3K27M (n=6) versus H3WT [n=4, two pediatric GBM (SF188, SJGBM2), one adult GBM (TS543), and an IDH1 R123H mutant (TS603) cell line] patient-derived cell lines (**Fig 1C**). As internal controls all H3K27M vs H3WT cells showed global reduction of H3K27me3, expression of H3K27M, and higher H3K27ac levels (**Fig 1B-C**). Both H3.3 and H3.1K27M versus H3WT cells showed higher H3K27Lac levels in IUE cells (**Fig 1B**). Similarly, H3K27Lac levels were significantly (p=0.0003) higher in H3K27M versus H3WT patient-derived cells (**Fig 1C-D**). Both H3K27Lac and H3K27ac are histone lysine-acylation marks.^44^ We tested if H3K27Crotonylation (Cr), another acylation mark, is similarly elevated in H3K27M cells. In contrast to H3K27Lac, H3K27Cr levels were not different between H3K27M and H3WT cells (**Fig 1C-D**).

We next sought to determine the effect of suppressing H3K27M mutation on H3K27Lac. We assessed two independent patient-derived isogenic cell lines with either knockdown (DIPG007 cells with or without shRNA H3-3A knockdown) or CRISPR correction (M corrected to K in BT245 cells^18^, henceforth referred to as CRISPR) (**Fig 1E**). Western blotting confirmed reduction of mutant H3K27M and H3K27ac with restoration of global H3K27me3 levels in both isogenic pairs (**Fig. 1F**). This was accompanied by a global reduction in both H3K27Lac and H3K27ac. Other marks including H3K14Lac, H3K9Lac, or H3K18Lac did not exhibit a consistent phenotype (**Fig. 1F**). Similar experiments in H3G34R/V mutant high grade glioma cells with CRISPR corrected isogenic pairs (**Fig S1F-G**). did not show consistent global changes in multiple H3KLac (H3K27Lac, H3K18Lac, H3K14Lac, H3K9Lac), H4K12Lac, and H2AK11Lac marks (**Fig S1H-I**).

Our data suggests that amongst the ten lactyl marks we profiled in two different histone mutant glioma models, H3K27Lac is the only lactylation mark that is globally higher in H3.3/H3.1K27M versus H3WT or H3G34 mutant cells isogenic mNSC, and H3K27M versus H3WT adult and pediatric GBM, and IDH1 mutant patient-derived cell lines.

### Suppressing H3K27M lowers intergenic H3K27Lac enrichment at ERV elements

We assessed changes in genomic distribution of H3K27Lac by performing chromatin immunoprecipitation sequencing (ChIP-seq) in isogenic cells with or without H3K27M suppression. We noted overall lowered global genomic reduction of H3K27Lac in both DIPG007 and BT245 cells compared to their corresponding H3K27M suppressed counterparts (**Fig 2A**). H3K27ac is pervasively distributed through the genome, including at promoters, enhancers, and intergenic regions including endoretroviral (ERV) elements in H3K27M cells.^21^ This prompted us to evaluate H3K27Lac distribution throughout the genome in our isogenic cells.

**Figure 2.**
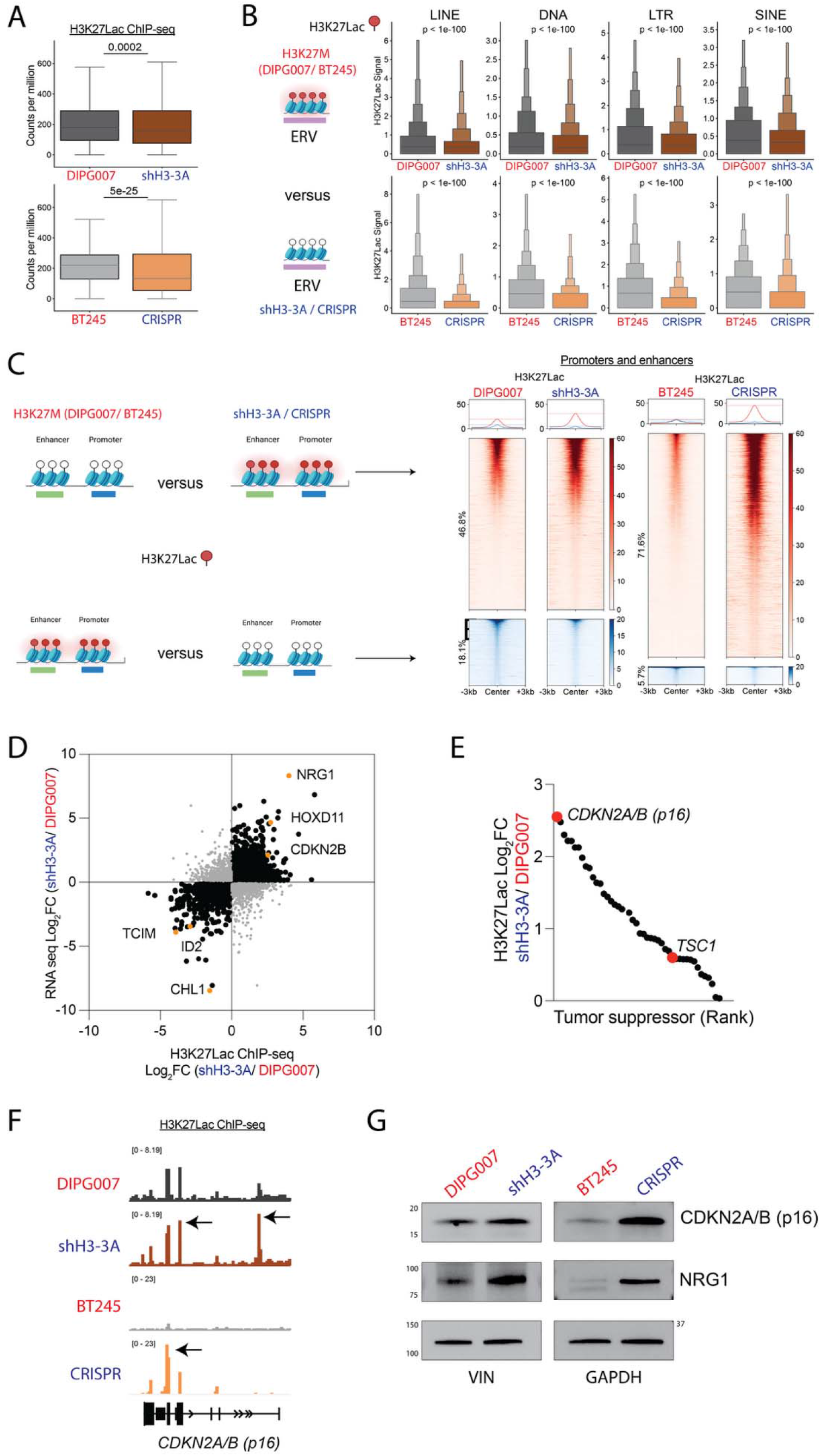
Suppressing H3K27M lowers intergenic H3K27Lac enrichment at ERV elements. - **(A)** Box-and-whisker plots showing normalized H3K27Lac read counts (counts per million) distribution in DIPG007 versus shH3-3A (top), and BT245 versus CRISPR (bottom) cells. Data analyzed using Mann-Whitney U test. Boxes show the interquartile range (IQR), with median lines; whiskers extend to the most extreme values within 1.5 × IQR of the box, with P-values shown above each comparison.
- **(B)** <u>Left</u>: Schema summarizing H3K27Lac occupancy in ERV regions in DIPG007 and BT245 (top) and shH3-3A/ CRISPR (bottom) cells. <u>Right</u>: Boxen plots showing the quartile distribution of H3K27Lac enrichment and median enrichment score across LINE, DNA, LTR, and SINE classes of transposable elements in DIPG007 versus shH3-3A (top), and BT245 versus CRISPR (bottom) cells. Central boxes span the 25th–75th percentiles, with progressively narrower boxes representing successive halving of each tail. Statistical significance was determined using a two-sided Mann–Whitney U tests, with P-values displayed above each comparison.
- **(C)** <u>Left</u>: Schema summarizing H3K27Lac occupancy in promoter and enhancer regions in DIPG007, BT245, shH3-3A, and CRISPR cells. <u>Right</u>: Heatmaps showing H3K27Lac ChIP-seq enrichment centered on peak (±3 kb) at promoters and enhancers for DIPG007 versus shH3-3A (left), and BT245 versus CRISPR (right). Individual peaks are displayed as rows and ordered by signal intensity. Metaplots above each heatmap depict the mean signal across all peaks. Numbers on the side of heatmaps indicate percent change within each comparison; peaks with similar signal are not shown (methods).
- **(D)** Scatter plot showing H3K27Lac ChIP-seq occupancy (Log_2_FC; x-axis) and differential gene expression (RNA-seq, Log_2_FC; y-axis) in shH3-3A to DIPG007 cells. Genes in the upper right quadrant exhibit upregulation in H3K27Lac occupancy and transcript expression. Genes in the lower left quadrant exhibit downregulation in H3K27Lac occupancy and transcript expression.
- **(E)** Ranked tumor suppressor by H3K27Lac enrichment in shH3-3A versus DIPG007 cells. Each point denotes a tumor suppressor gene, with the y-axis indicating the log_2_FC in H3K27Lac enrichment at the corresponding genomic locus.
- **(F)** Representative H3K27Lac ChIP tracks at the *CDKN2A/B* locus in DIPG007 versus shH3-3A (top), and BT245 versus CRISPR (bottom) H3K27M cells.
- **(G)** Representative Western blots of p16, NRG1, vinculin, and GAPDH in DIPG007 versus shH3-3A (left), and BT245 versus CRISPR (right) H3K27M cells. Figures in panel B and C were generated using biorender.

**Figure S2.**
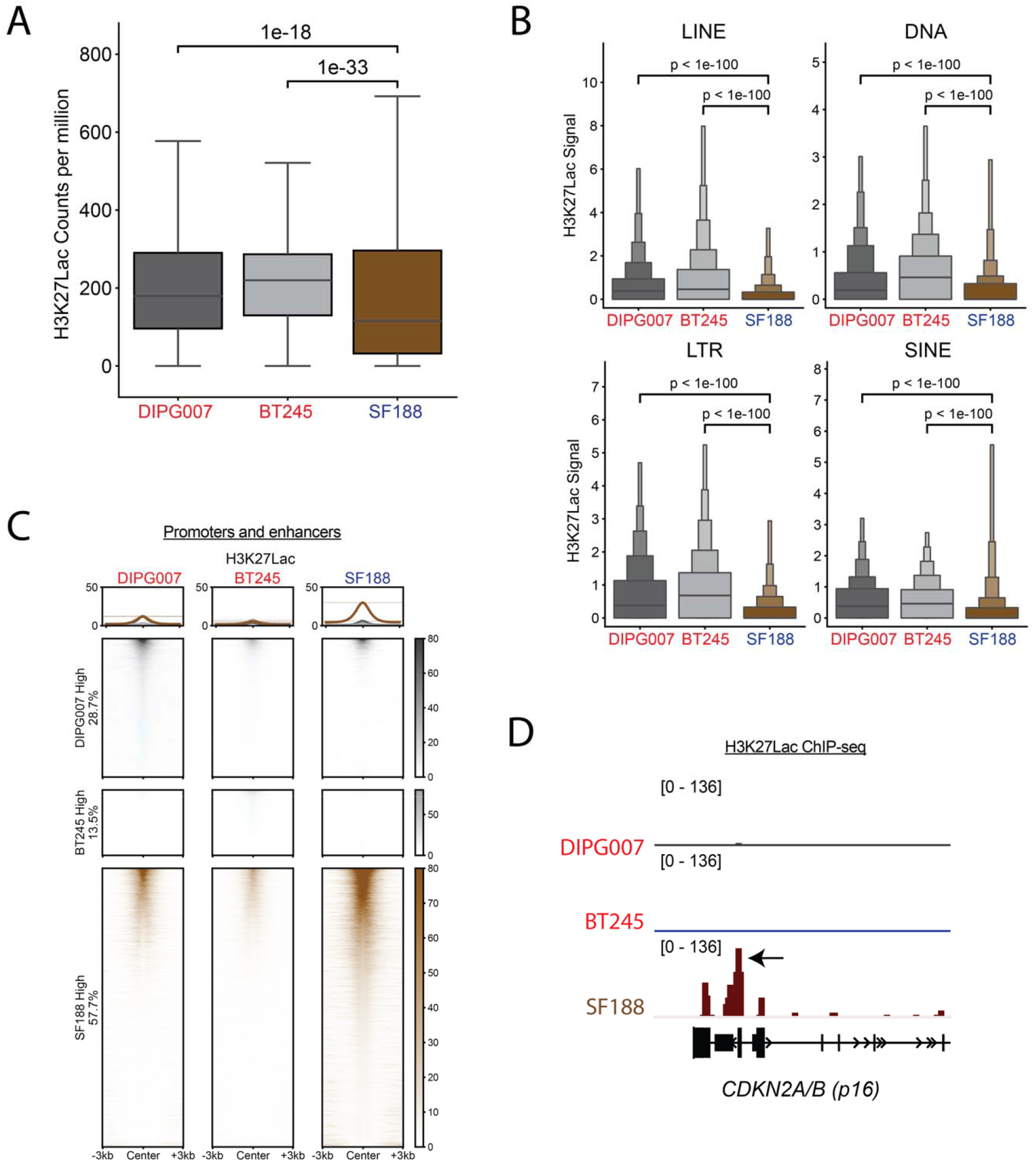
Suppressing H3K27M lowers intergenic H3K27Lac enrichment at ERV elements (related to figure 2) - **(A)** Box-and-whisker plots showing normalized H3K27Lac read counts (counts per million) distribution in H3K27M (DIPG007, left; BT245, middle) versus H3WT SF188 (right) cells. Data analyzed using Mann-Whitney U test. Boxes show the interquartile range (IQR), with median lines; whiskers extend to the most extreme values within 1.5 × IQR of the box, with P-values shown above each comparison.
- **(B)** Boxen plots showing the quartile distribution of H3K27Lac enrichment and median enrichment score across LINE, DNA, LTR, and SINE classes of transposable elements in in DIPG007, BT245, and SF188 (left to right). Central boxes span the 25th–75th percentiles, with progressively narrower boxes representing successive halving of each tail. Statistical significance was determined using a two-sided Mann–Whitney U tests, with P-values displayed above each comparison.
- **(C)** Heatmaps showing H3K27Lac ChIP-seq enrichment centered on peak (±3 kb) at promoters and enhancers for DIPG007, BT245, and SF188 (left to right). Individual peaks are displayed as rows and ordered by signal intensity. Metaplots above each heatmap depict the mean signal across all peaks. Numbers on the side of heatmaps indicate percent change within each comparison; peaks with similar signal are not shown (methods).
- **(D)** Representative H3K27Lac ChIP tracks at the *CDKN2B* locus in DIPG007 (top), BT245 (middle), and SF188 (bottom) cells.

Suppression of H3K27M resulted in lowering of H3K27Lac levels at intergenic regions including LINE, SINE, LTR, and DNA classes of ERV transposable repetitive elements in DMG cells (**Fig 2B).** We also examined H3K27M (DIPG007, BT245) versus H3WT (SF188) cells. These data showed a similar pattern. Overall genomic H3K27Lac levels were higher in H3K27M versus H3WT cells (**Fig S2A**). Moreover, H3K27Lac was higher in genomic LINE, SINE, LTR, and DNA ERV elements in both H3K27M cells versus H3WT cells (**Fig S2B**).

We next assessed H3K27Lac distribution at promoters and enhancers. Prior results have shown at active enhancers, H3K27M causes reduction of H3K27ac and lowered transcriptional expression of adjacent neurodevelopmental genes.^11,21^ H3K27Lac followed a similar overall pattern and was higher at promoter and enhancer regions in cells with H3K27M mutant suppression compared to their isogenic parental controls or H3WT SF188 cells (**Fig 2C, Fig S2C**).

To determine potential function relevance of changes at promoters and enhancers, we compared alterations in H3K27Lac with gene expression in DIPG007 cells with or without shH3-3A. We identified down-regulated genes that showed lower H3K27Lac deposition and *vice versa*. Amongst factors that showed low H3K27Lac, we noted downregulated expression of neurodevelopment related genes including *TCIM* (Transcriptional and Immune Response Regulator positive activator of the Wnt/beta-catenin signaling), *CHL1* (Cell Adhesion Molecule L1 that relates with brain development), and *ID2* (Inhibitor of DNA Binding 2 implicated in neuronal stem cell growth). On the other hand, top genes with both increased H3K27Lac and upregulation included *CDKN2B*, *HOXD11* (canonical PRC2 target), and *NRG1* (Neuregulin 1 encodes a CNS signaling protein related to differentiation) (**Fig 2D**). Prior studies have shown that *CDKN2A/B*, a tumor suppressor encoding p16, is specifically repressed by the H3K27M mutation.^20,45^ Targeting H3K27M led to increased H3K27Lac deposition at the *CDKN2A/B* locus and other tumor suppressor loci (**Fig 2E-F**). Similarly, H3K27Lac enrichment was higher at the *CDKN2A/B* locus in SF188 versus BT245 and DIPG007 cells (**Fig S2D**). We validated increased expression of key genes, including p16 and NRG1, at the protein level (**Fig 2G**). Overall, our findings suggest that suppressing H3K27M lowered overall H3K27Lac enrichment at intergenic regions including ERV elements. This was accompanied by increased H3K27Lac deposition and expression of genes related to neurodevelopment and tumor suppressors including p16.

### Lactate levels are heterogeneous in H3K27M DMGs

Lactate levels in tumors show both intertumoral and intratumor heterogeneity. For example, a study in adult glioblastoma (GBM) from eight patients show intratumor lactate ranges from 6.33 to 310 mM.^46^ Lactate levels can also vary within tumor microenvironments with hypoxic regions containing high concentrations of lactate.^47^ We sought to assess the impact of tumor microenvironmental lactate on H3K27Lac. We first assessed heterogeneity in lactate levels in H3K27M DMG models *in vitro* and *in vivo* using uniformly labeled [U-^13^C_6_] glucose tracing. We profiled nine patient, low passage cell lines derived from H3.3K27M (n=6) and H3.1K27M (n=3) patients (**Fig 3A**). Glucose derived lactate (containing three labelled carbons M+3) was detected in all cells but showed marked heterogeneous levels with no notable differences between H3.3K27M and H3.1K27M cells. Similarly, [U-^13^C_6_] *in vivo* glucose tracing was performed in two independent orthotopic H3K27M DMG animal models. Amongst these, DIPG007 versus PPK orthotopic implanted animals showed a greater portion of glucose-derived lactate (**Fig 3B**). To see if patient H3K27M DMGs show similar heterogeneity, we used magnetic resonance spectroscopy (MRS) to assess intertumoral lactate levels in H3K27M DMG including ten H3.3K27M and two H3.1K27M patients *in vivo*. Similar to our experimental models, we noted variability in tumor lactate levels amongst H3K27M DMG patients without major differences between H3.3K27M and H3.1K27M patients (**Fig 3C-F**). From our analysis, lactate levels are heterogeneous amongst H3K27M DMGs and prompted us to determine the effect of exogenously increasing lactate concentrations on H3K27Lac in H3K27M DMGs.

**Figure 3.**
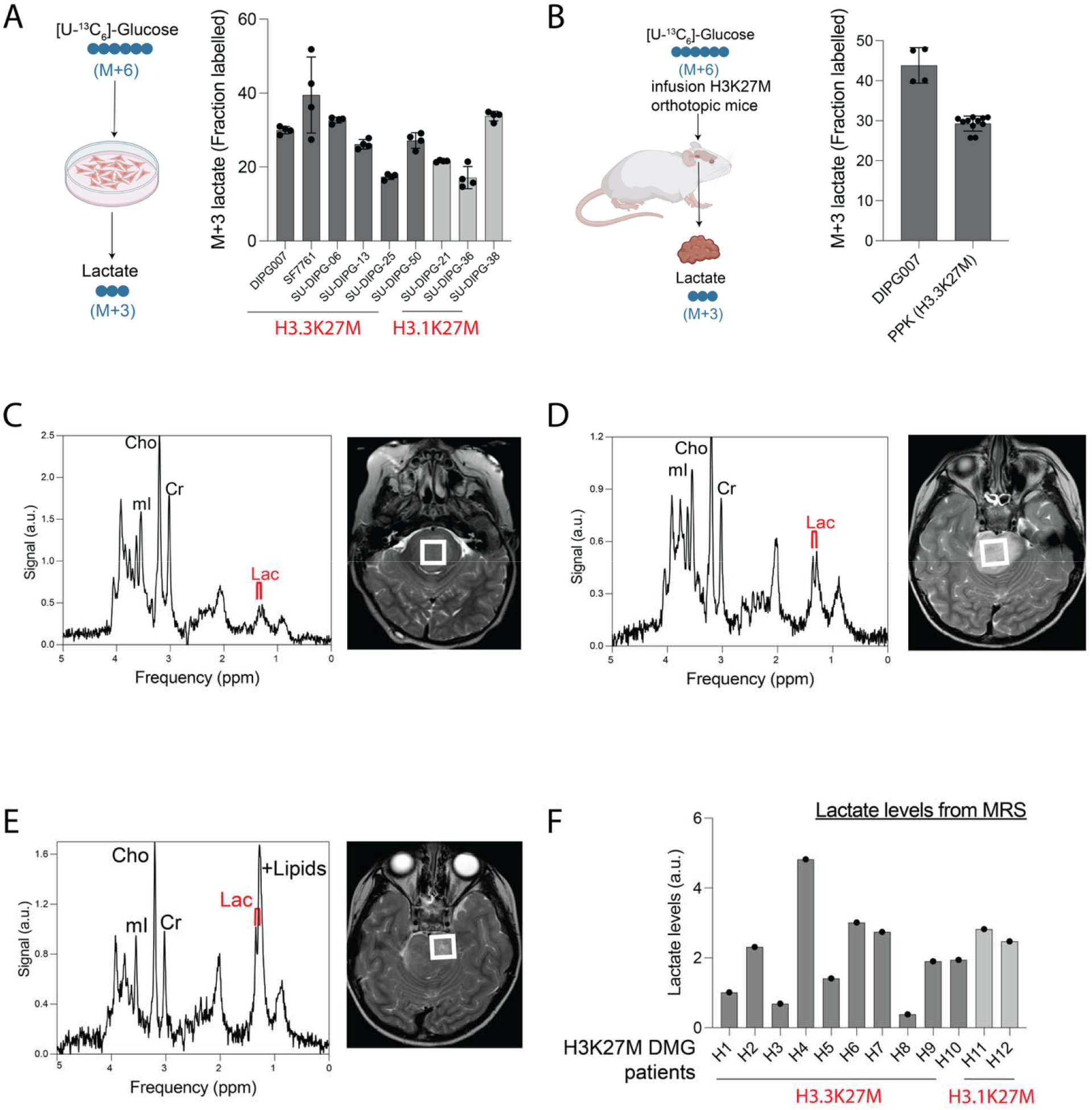
Lactate levels are heterogeneous in H3K27M DMGs. - **(A)** <u>Left</u>: Schematic showing uniformly labeled [U-^13^C_6_]-glucose (with 6 labelled carbons; M+6) tracing to assess glucose-derived lactate (with 3 labelled carbons; M+3) *in vitro*. <u>Right</u>: Bar chart illustrating fraction of labeled M+3 lactate in H3.3K27M (n=6) and H3.1K27M (n=3) cell lines. Data shown as mean ± SD.
- **(B)** <u>Left</u>: Schematic showing uniformly labeled [U-^13^C_6_]-glucose with 6 labelled carbons; M+6) tracing to assess glucose-derived lactate (with 3 labelled carbons; M+3) *in vivo*. H3K27M-bearing mice were infused with [U-^13^C_6_]-glucose (bolus dose (0.4 mg/g) followed by a continuous infusion (12 ng/g per min) for a total of 4 h. Tumors were rapidly extracted and analyzed using mass spectroscopy. <u>Right</u>: Bar chart illustrating fraction of labeled M+3 lactate in H3K27M tumor tissues bearing H3.3K27M DIPG007 (DIPG007, n=4) or PPK (n=11) cells. Data shown as mean ± SD.
- **(C)** –(**E**) <u>Left</u>: Representative magnetic resonance spectroscopy (MRS) spectrum with metabolite resonances labeled for myo-inositol (mI), choline (Cho), creatine (Cr), lactate (Lac), and lipids from three H3K27M DMG patients. Signal intensity is shown on the y-axis in arbitrary units (a.u.); x- axis shows frequency as a function of chemical shift (ppm). <u>Right</u>: axial T2-weighted MRI illustrating the tumor area selected for MRS acquisition (white square).
- **(F)** Bar chart showing MRS quantified lactate levels (y-axis, a.u.) in H3.3K27M (n=10) and H3.1K27M (n=2) DMG patients. Figures in panels A and B were generated using biorender.

### Exogenous lactate increases H3K27Lac to drive neuro-developmental pathways gene expression

To mimic high lactate DMGs *in vitro*, we treated H3K27M cell lines with lactate and tested if lactate supplementation could alter H3K27Lac levels. BT245 H3K27M cells treated with increasing concentrations (0-20mM) of lactate showed a dose-dependent increase in H3K27Lac (**Fig 4A**). No differences were noted in H3K27ac levels (**Fig 4A**). DIPG007 and H3.3K27M mNSCs also showed a dose-dependent increase in H3K27Lac levels (**Fig 4B, Fig S3A**). Because lactate is secreted by tumor cells, we treated DIPG007 and BT245 cells treated with their respective conditioned media. Like our exogenous lactate experiments, conditioned media also increased H3K27Lac levels (**Fig 4C**).

**Figure 4.**
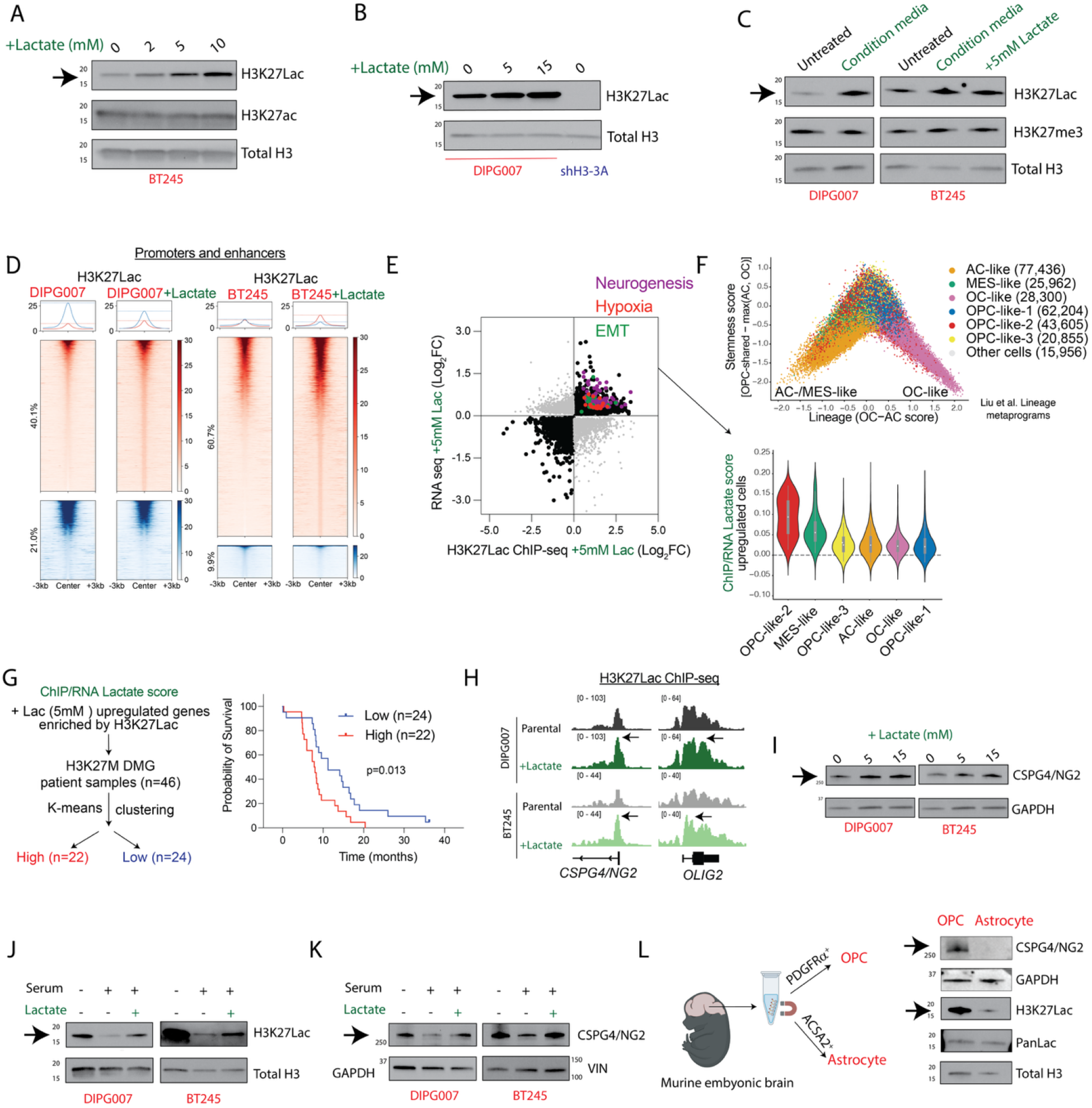
Exogenous lactate increases H3K27Lac to drive neuro-developmental pathways gene expression. - **(A)** Representative Western blots of H3K27Lac (arrow), H3K27ac and total H3 in H3K27M BT245 cells supplemented with increasing concentration of lactate (left to right: 0, 2, 5, 10, 20m M) for 24 h.
- **(B)** Representative Western blots of H3K27Lac (arrow) and total H3 in DIPG007 cells supplemented with 0, 5, or 15 mM lactate for 24 h; shH3-3A cells included as control.
- **(C)** Representative Western blots of H3K27Lac (arrow), H3K27me3, and total H3 in DIPG007 and BT245 in the following conditions (left to right: DIPG007 untreated, DIPG007 tumor conditioned media-treated, BT245 untreated, BT245 tumor conditioned media-treated, and BT245 with 5mM exogenous lactate supplementation) for 24h.
- **(D)** Heatmaps showing H3K27Lac ChIP-seq enrichment centered on peaks (±3 kb) at promoters and enhancers for DIPG007 versus DIPG007 + lactate, or BT245 versus BT245 + lactate. Individual peaks are displayed as rows and ordered by signal intensity. Red heatmaps denote genomic regions with higher H3K27Lac occupancy, blue heatmaps denote genomic regions with lower H3K27Lac occupancy, and the corresponding color scales are displayed on the right. Metaplots above each heatmap depict the mean signal across all peaks. Numbers on the side of heatmaps indicate percent change within each comparison; peaks with similar signal are not shown (methods). Cells were treated with 5mM lactate for 48h.
- **(E)** Scatter plot showing H3K27Lac ChIP-seq occupancy (Log_2_FC; x-axis) versus differential gene expression (RNA-seq log2FC; y-axis) in isogeneic DIPG007 + 5mM lactate relative to untreated DIPG007 cells. Genes in the upper right quadrant exhibit increases in H3K27Lac occupancy and transcript expression. Genes in the lower left quadrant exhibit decreases in H3K27Lac occupancy and transcript expression. Genes mapping to the “neurogenesis,” “hypoxia,” and “EMT (epithelial to mesenchymal transition)” pathways are color coded (GSEA).
- **(F)** Top: Wishbone-plot analysis (x-axis, lineage score; y-axis stemness score) of published H3K27M DMG single-cell RNA-seq data ^24,77^ showing tumor cell state metaprograms: Astrocyte (AC)-like, n=77,436 cells; mesenchymal (MES)-like, n=25,962 cells; oligodendrocyte (OC)-like, n=28,300 cells; oligodendrocyte precursor (OPC)-like 1, n=62,204 cells; OPC-like 2, n=43,605 cells; and OPC-like 3, n=20,855 cells. Bottom: Violin plots showing the projected distribution of cells (wishbone plot on top) with genes that showed increased H3K27Lac enrichment and upregulated expression in DIPG007 lactate treated cells (taken from fig 4E, upper right quadrant). The X-axis indicated cell states. The width of each violin represents the density of observations at each value. The white dot denotes the median, and the gray box indicates the interquartile range (IQR), with whiskers representing the distribution range.
- **(G)** Genes that showed H3K27Lac enrichment and upregulated expression in DIPG007 lactate treated cells (upper right quadrant, from E) were derived from published H3K27M clinical data sets.^48^ Gene expression data were stratified into two H3K27M patient groups by unbiased K-means clustering into high (n=22, red) versus low (n=24, blue) groups. Kaplan-Meier survival analysis (with 95% C.I.) of published patient overall survival was performed between the two groups. P value is indicated.
- **(H)** Representative H3K27Lac ChIP tracks at *CSPG*4/NG2 and *OLIG2 loci* in DIPG007 versus DIPG007+lactate, and BT245 versus BT245+lactate isogenic conditions. Cells were treated with 5mM lactate for 48h.
- **(I)** Representative Western blots of CSPG4/NG2 (arrow) and GAPDH in DIPG007 and BT245 supplemented with 0, 5, or 15mM lactate for 48 h.
- **(J)** Representative Western blots of H3K27Lac (arrow) and total H3 in DIPG007 (left) and BT245 (right) in the following condition: control, (+) 10% fetal bovine serum (FBS) for 2 weeks, and (+) 10% fetal bovine serum (FBS) for 2 weeks with exogenous lactate supplementation for 24 h before cell collection.
- **(K)** Representative Western blots of CSPG4/NG2 (arrow), GAPDH, and vinculin (VIN) in DIPG007 (left) and BT245 (right) in the following condition: control, (+) 10% fetal bovine serum (FBS) for 2 weeks, and (+) 10% fetal bovine serum (FBS) for 2 weeks with exogenous lactate supplementation for 24 h before cell collection.
- **(L)** <u>Left</u>: Scheme showing isolation of primary murine OPC cells with CD140a/PDGFRα microbeads, and or astrocytes with ACSA2 microbeads were isolated from dissociated murine E18 brain tissues. <u>Right</u>: Representative Western blots of CSPG4/NG2 (arrow), GAPDH, H3K27Lac (arrow), Pan lysine (K) Lac, and total H3 in both cell types. Figures in panel L were generated using biorender.

**Figure S3.**
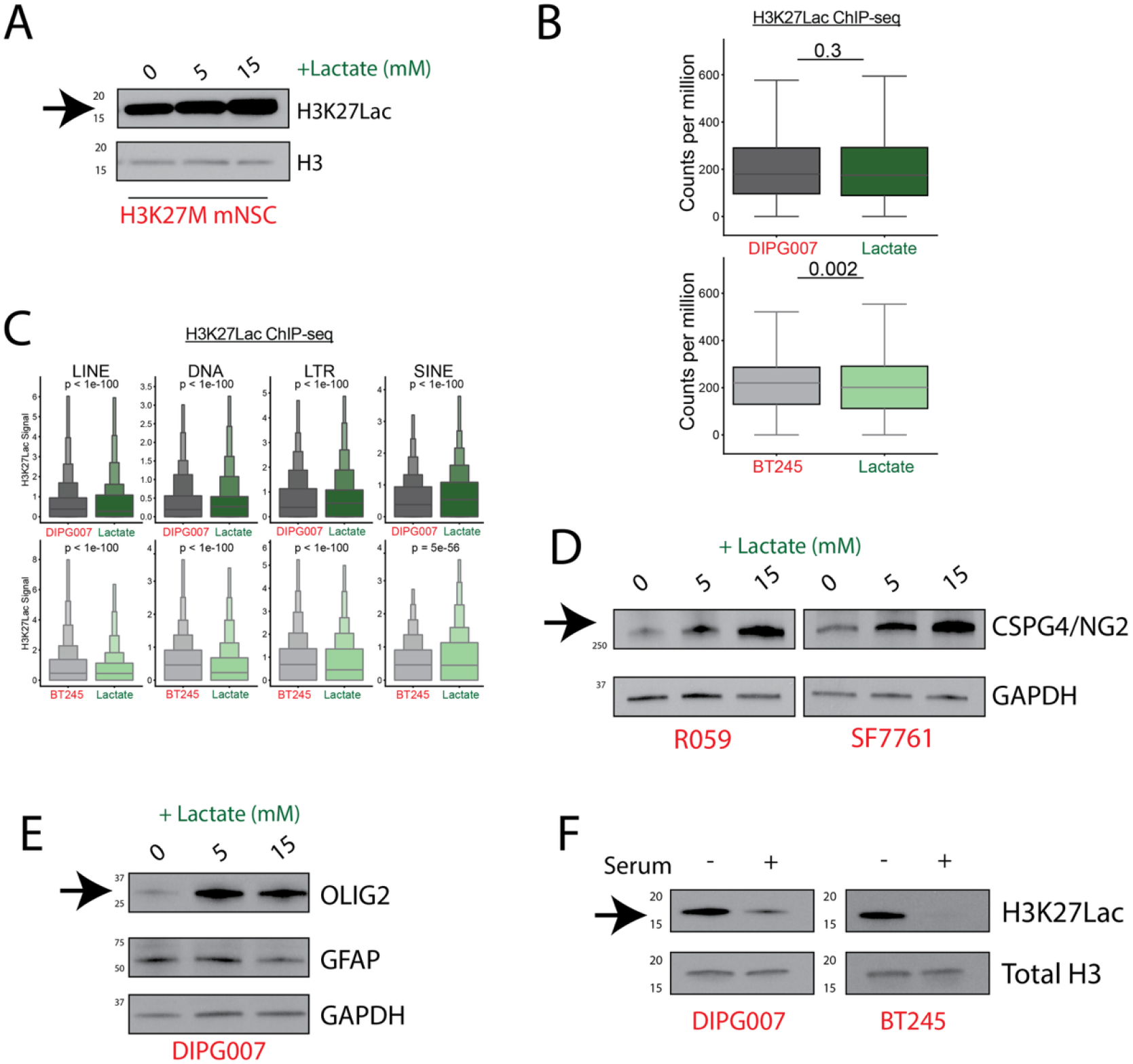
Exogenous lactate increases H3K27Lac to drive neuro-developmental pathways gene expression (related to figure 4). - **(A)** Representative western blots of H3K27Lac (arrow) and total H3 in H3K27M mNSCs supplemented with increasing concentration of lactate (left to right: 0, 5, 15mM) for 24 h.
- **(B)** Box-and-whisker plots showing normalized H3K27Lac read counts (counts per million) distribution in DIPG007 versus DIPG007 + lactate (top), and BT245 versus BT245 + lactate isogenic cells (bottom). Data analyzed using Mann-Whitney U test. Boxes show the interquartile range (IQR), with median lines; whiskers extend to the most extreme values within 1.5 × IQR of the box, with P-values shown above each comparison. Cells were treated with 5mM lactate for 48h.
- **(C)** Boxen plots showing the quartile distribution of H3K27Lac enrichment and median enrichment score across LINE, DNA, LTR, and SINE classes of transposable elements in DIPG007 versus DIPG007 + lactate (top), and BT245 versus BT245 + lactate (bottom) isogenic cells. Central boxes span the 25th– 75th percentiles, with progressively narrower boxes representing successive halvings of each tail. Statistical significance was determined using a two-sided Mann–Whitney U tests, with P-values displayed above each comparison. Cells were treated with 5mM lactate for 48h.
- **(D)** Representative Western blots of CSPG4/NG2 (arrow) and GAPDH in R059 and SF7761 H3K27M cells supplemented with 0, 5, or 15mM lactate for 48 h.
- **(E)** Representative Western blots of OLIG2 (arrow), GFAP, and GAPDH in DIPG007 H3K27M cells supplemented with 0, 5, or 15mM lactate for 48 h.
- **(F)** Representative Western blots of H3K27Lac (arrow) and total H3 in DIPG007 (right) or BT245 (left) H3K27M cells cultured with (+) or without (-) 10% fetal bovine serum (FBS) for 2 weeks.

We performed ChIP-seq to determine changes in genomic distribution of H3K27Lac on exogenous lactate addition (5mM for 48 h). Global genomic H3K27Lac including at intergenic regions did not show a consistent pattern of difference in DIPG007 and BT245 cells on lactate addition (**Fig S3B-C**). However, we noted increased H3K27Lac enrichment at several enhancer and promoter regions in both cell lines (**Fig 4D**). To determine functional significance, we compared changes in H3K27Lac genomic distribution with gene expression in DIPG007 cells on lactate addition. Genes with increased H3K27Lac enrichment and increased gene expression corresponded to neurogenesis, hypoxia, and EMT-related signatures (**Fig 4E**). H3K27M DMGs are heterogeneous and contain different tumor states including cells within OPC--like (groups 1, 2 and 3), OC-like, AC-like, and MES-like states.^23–29^ We assessed if lactate-driven genes (upregulated with exogenous lactate and enriched with H3K27Lac) corresponded to these cell states. Amongst these, lactate-driven genes showed most overlap with OPC-like-2 and MES-like cell states (**Fig 4F**). Because differentiation-related states in DMGs can relate with prognosis,^23–29^ we assessed the potential clinical significance of these lactate-driven genes. We used K-means clustering of lactate-driven genes to stratify H3K27M DMG (n=46) patients into high versus low expression groups.^48^ High-versus-low expression of lactate-driven genes showed worse overall survival outcomes in H3K27M DMGs (**Fig 4G**).

Amongst lactate-driven genes, we noted that key neuronal stem cells/ OPC-like markers including *CSPG4/ NG2* and *OLIG2* showed increased genomic H3K27Lac enrichment on addition of 5mM lactate in DIPG007 and BT245 cells (**Fig 4H**). NG2 and OLIG2 protein levels were also upregulated on treating H3K27M cells with increasing concentrations of lactate (**Fig 4I, Fig S3D-E**). H3K27M tumor cell states are plastic, and OPC-like cells can be forcibly reprogrammed to AC-like H3K27M cells by culturing them with serum.^23,49^ Addition of serum to DIPG007 and BT245 cells reduced H3K27Lac levels (**Fig 4J; Fig S3F**). This phenotype was rescued by the addition of 5mM lactate (**Fig 4J**). Functionally, serum treatment reduced NG2 levels, which was reversed by exogenous lactate (**Fig 4K**). These data suggest the hypothesis that exogenous lactate promotes a more stem cell/OPC-like state. To test this, we isolated OPCs and ACs from wildtype embryonic mice. OPCs versus ACs showed higher levels of NG2 and H3K27Lac, supporting this hypothesis (**Fig 4L**). Thus, lactate addition increased H3K27Lac enrichment and expression at key genes related to neuronal stem cells/ OPC-like states and high expression of lactate-driven genes corresponded to H3K27M DMGs with overall poor outcome.

### Lactate increases H3K27Lac to activate mesenchymal related genes

Because we noted that lactate-driven genes also related with EMT and hypoxia gene signatures (**Fig 4E-F**), we focused our efforts on understanding the effect of lactate on H3K27M DMG mesenchymal-like states. Lactate addition to DIPG007 and BT245 increased genomic enrichment of H3K27Lac at key hypoxia and epithelial-to-mesenchymal (EMT) transition factors including *GALK1* and *TGFB1* respectively (**Fig 5A**). Moreover, lactate treatment increased expression of several EMT and hypoxia-related gene signatures in both cells (**Fig S4A**). We also noted a lactate driven, dose-dependent increased expression of several of these genes, including GALK1 and TGFB1 in DIPG007 cells (**Fig 5B**).

**Figure 5.**
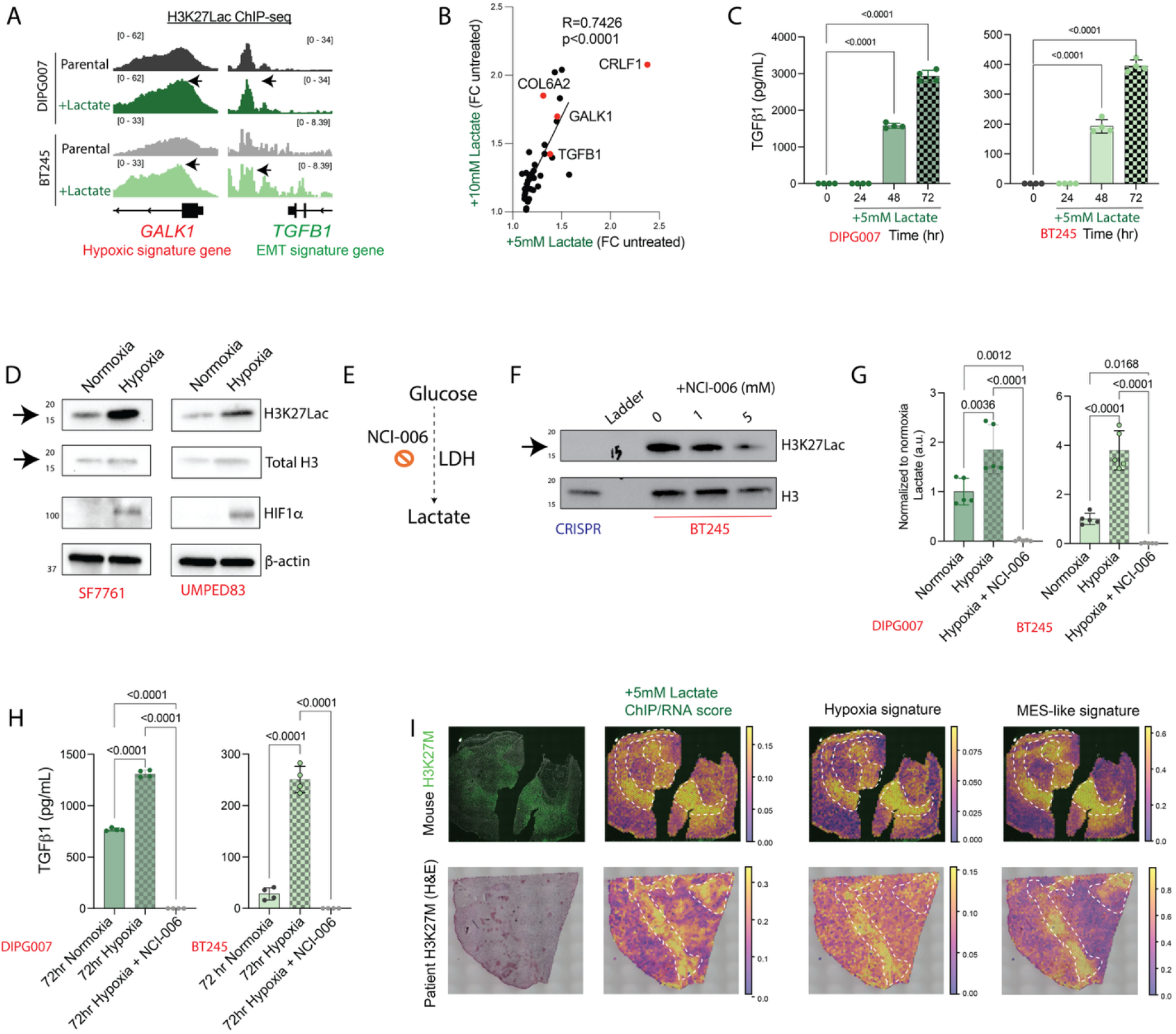
Lactate increases H3K27Lac to activate mesenchymal related genes. - **(A)** Representative H3K27Lac ChIP tracks at *GALK1* and *TGFB1 loci* in DIPG007 versus DIPG007+lactate, and BT245 versus BT245+lactate isogenic conditions. Cells were treated with 5mM lactate for 48h.
- **(B)** Scatter plot showing normalized RNA-seq expression values in fold change for DIPG007 + 5mM lactate relative to untreated (x-axis, n=3 each) and DIPG007 + 10mM lactate relative to untreated (y-axis, n=3 each). Each point denotes one gene, and the solid line denotes the fitted linear regression. Data were analyzed by Pearson correlation using 95% C.I. Pearson’s R and P values are indicated.
- **(C)** Bar graphs showing TGFβ1 concentration (pg/mL, y-axis) measured by ELISA in vehicle- or lactate-treated (for 24, 48, or 72 h) DIPG007 (left) and BT245 (right) media. Data shown as mean ± SD and analyzed by one-way ANOVA with a Tukey’s multiple comparisons using 95% C.I. P values are indicated.
- **(D)** Representative Western blots of H3K27Lac (arrow), total H3, HIF1α, and β-actin in SF7761 (left) and UMPED83 (right) cells cultured in normoxic versus hypoxic conditions for 48h.
- **(E)** Diagram showing lactate generation from glucose by the enzyme lactate dehydrogenase (LDH). NCI-006 is an LDH specific inhibitor.
- **(F)** Representative Western blots of H3K27Lac (arrow) and total H3 in BT245 parental H3K27M cells treated with 0, 1, or 5 mM of NCI-006. Isogenic CRISPR cells were used as control.
- **(G)** Bar graphs showing extracellular lactate levels (mM) measured via a colorimetric assay in DIPG007 (left) and BT245 (right) cells cultured in normoxia, hypoxia, or hypoxia treated with NCI-006 (2.5mM) for 72-hr.
- **(H)** Bar graphs showing TGFβ1 concentration (pg/mL, y-axis) measured by ELISA in DIPG007 (left) and BT245 (right) cells cultured in normoxia (base line), hypoxia, or hypoxia treated with NCI-006 (2.5mM) for 48 or 72 h. Data shown as mean ± SD and analyzed by two-way ANOVA with adjusted p values shown after Šídák’s multiple comparisons using 95% C.I. P values are indicated.
- **(I)** Representative spatial maps from tissue samples (top row: mouse H3K27M tumors; bottom row: patient H3K27M sample; data derived from Colot et al. 2026).
  – The first column shows H3K27M stained murine H3K27M tumor tissue (top) or hematoxylin and eosin (H&E) stained patient H3K27M DMG (bottom).
  – The second column shows spatial enrichment of lactate driven ChIP/RNA score (genes with H3K27Lac enrichment and upregulated expression in DIPG007 cells treated with 5mM lactate from main figure 4E-G).
  – The third and fourth columns show spatial enrichment of hypoxia and MES-like gene signatures. The dashed white outlines delineate the area of interest. Color intensity represents the magnitude of the corresponding molecular score, with scales shown next to each panel.

**Figure S4.**
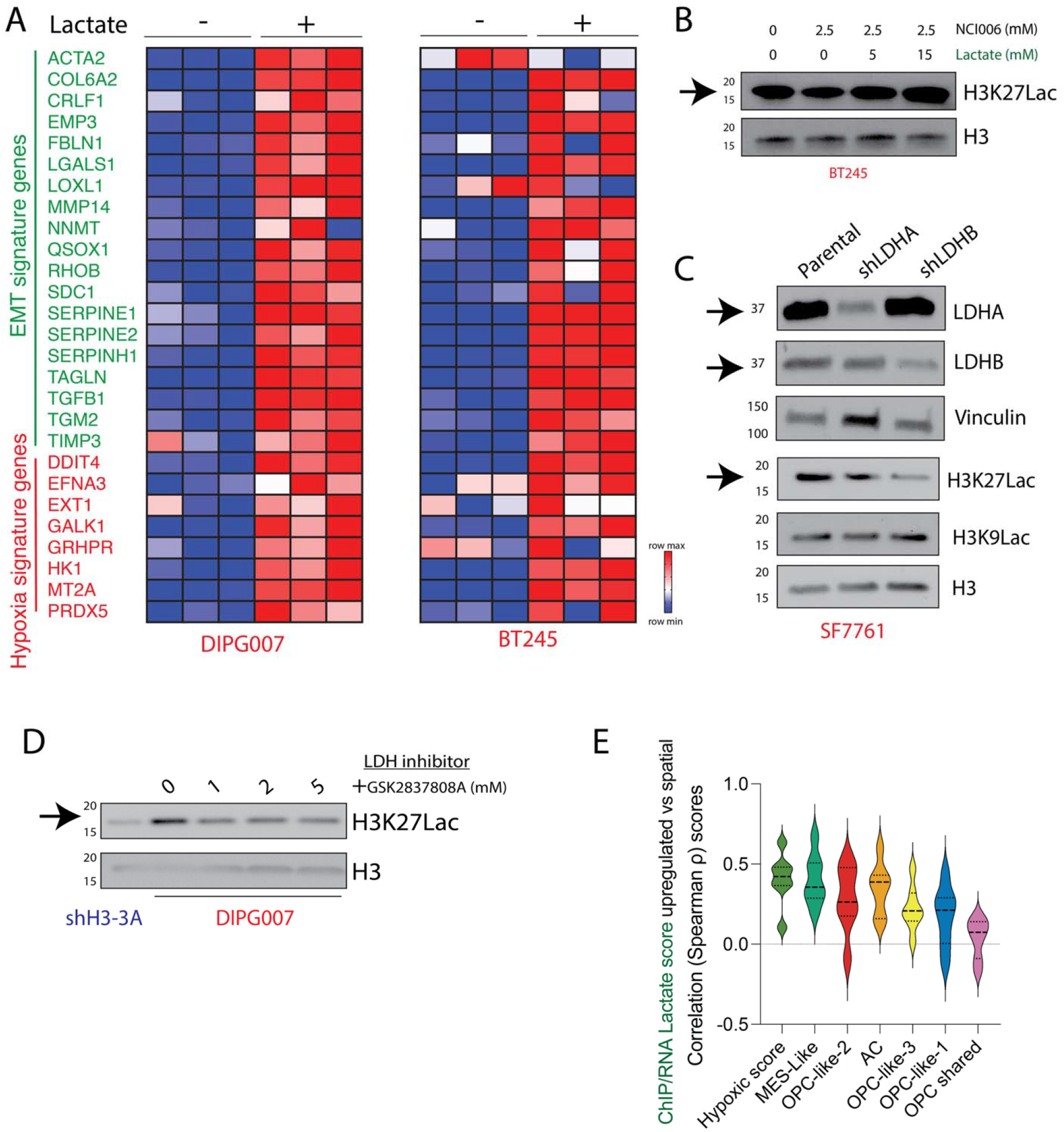
Lactate increases H3K27Lac to activate mesenchymal related genes (related to figure 5). - **(A)** Heatmaps representing gene expression values of key EMT (green) and hypoxia (red) signature gene in DIPG007 (left) and BT245 (right) with or without 48-hr 10mM lactate supplementation (n=3, all conditions).
- **(B)** Representative Western blots of H3K27Lac (arrow) and total H3 in BT245 treated with 0 or 2.5uM NCI-006 with or without 0, 5, or 15mM lactate addback for 48h.
- **(C)** Representative Western blots of LDHA (arrow), LDHB (arrow), vinculin, H3K27Lac (arrow), H3K9Lac, and total H3 in SF7761 with or without shRNA-mediated stable knockdown of LDHA and LDHB.
- **(D)** Representative Western blots of H3K27Lac (arrow) and total H3 in DIPG007 treated with 0, 1, 2, or 5 mM of LDHA inhibitor GSK2837808A for 48h. shH3-3A cells used as control.
- **(E)** Violin plots showing correlation of integrated upregulated ChIP/RNA Spearman ρ score with spatial transcriptomics of indicated cell states (x-axis) after 5mM lactate treatment. The width of each violin represents the density of observations at each value. The black dashed horizontal lines within each violin indicate the 25th, 50th (median), and 75th percentiles, respectively. The height of each violin indicates the range of distribution.

Because TGFβ1 is a critical EMT and MES-regulating factor, we assessed secreted TGFβ1 protein levels on lactate treatment using ELISA. Addition of 5mM lactate was sufficient to increase TGFβ1 levels to greater than 200-fold by 48hrs in both DIPG007 and BT245 cells (**Fig 5C**). Since hypoxic tumor cells produced lactate, we assessed the effects of hypoxia on H3K27Lac and TGFβ1 production in H3K27M patient-derived cells. Incubating DIPG007 and BT245 cells in hypoxia increased lactate accompanied by elevated H3K27Lac levels (**Fig 5D-G**). To determine if this effect was dependent on lactate production, we treated cells with the lactate dehydrogenase (LDH) inhibitor, NCI-006 (**Fig 5E-G**).^50^ NCI-006 lowered both lactate and H3K27Lac levels (**Fig 5F-G**). H3K27Lac reduction on NCI-006 treated cells was reversed by competing out inhibitory effects with excess lactate (**Fig S4B**). As controls, genetic knockdown of LDHA and LDHB using shRNA also reduced H3K27Lac levels (**Fig S4C**). Similarly, another pharmacological inhibitor of LDHA (GSK2837808A) reduced H3K27Lac in a dose-dependent manner (**Fig S4D**). Increased lactate production and elevated H3K27Lac in hypoxia (**Fig 5D, G**) was associated with an increase in TGFβ1 secretion (**Fig 5H**). Notably, treatment with NCI-006 abrogated TGFβ1 production in hypoxia (**Fig 5H**).

We validated our findings using previously published spatial transcriptomic data derived from H3K27M patient samples (n=3) and mouse H3K27M DMG tumor tissues (n=4).^28^ We determined hypoxic and MES-like signatures and compared them to lactate-driven genes (**Fig 4E-G**). We noted a positive correlation between spatial distribution of lactate-driven genes with hypoxia and MES-like signatures (**Fig 5I, Fig S4E**). Our data demonstrates that exogenous lactate enriches genomic H3K27Lac, accompanied by increased expression of key hypoxic, EMT driving and MES-like genes including TGFB1. These key findings were corroborated in H3K27M DMG tumor single-cell and spatial transcriptomics data.

### CBP/p300 inhibition simultaneously lowers H3K27Lac and H3K27ac and decreases their enrichment at ERV elements

Histone lysine lactylation and acetylation are regulated by similar enzymes.^39^ For example, the enzyme ACSS2 (Acyl-CoA Synthetase Short Chain Family Member 2) functions as both lactyl-CoA and acetyl-CoA synthetase to increase H3K18Lac and H3K18ac levels in GBM cells (**Fig S5A**).^51^ Similarly, the enzymes CBP/p300 function as both lactyl- and acetyl-transferase to enhance both H3K18Lac and H3K18ac levels (**Fig S5A**).^39,52,53^ We hypothesized that targeting these enzymes will provide a significant advantage of simultaneously suppressing both H3K27Lac and H3K27ac marks, enhancing potential therapeutic effectiveness through a dual approach.

Knockdown of ACSS2 using two independent shRNA reduced H3K27Lac levels (**Fig S5B-C**). A pharmacological inhibitor of ACSS2 (MTB-9655) reduced both H3K27Lac and H3K27ac in a dose-dependent manner in H3K27M cells (**Fig S5D**). Genetic suppression of CBP or p300 decreased H3K27Lac as well as H3K27ac levels in R059 H3K27M cells (**Fig 6A-B**). We took advantage of a CBP/p300 PROTAC degrader, CBPD-409, which decreases CBP and p300 protein levels in nanomolar (nM) ranges.^54,55^ CBPD-409 reduced CBP and p300 protein levels accompanied by marked global reduction in H3K27Lac and H3K27ac in both BT245 and DIPG007 H3K27M cells (**Fig 6C-D**). H3K27M mutant levels did not change on CBPD-409 treatment (**Fig 6E**). We focused our efforts on targeting CBP/p300 with CBPD-409 because of its efficacy in reducing both H3K27Lac and H3K27ac in the nanomolar range.

**Figure 6.**
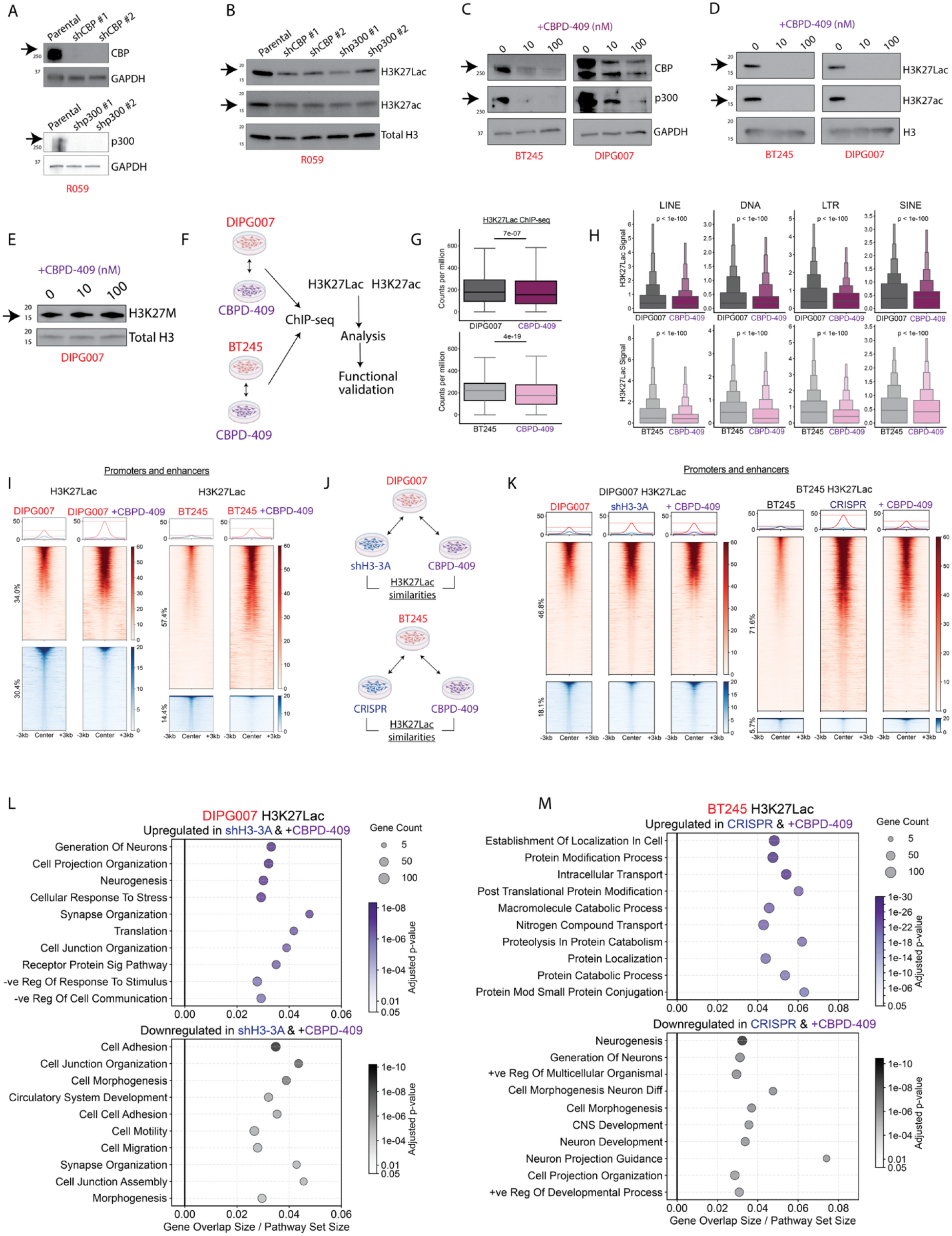
CBP/p300 inhibition simultaneously lowers H3K27Lac and H3K27ac and decreases their enrichment at ERV elements. - **(A)** Representative Western blots of CBP (arrow), p300 (arrow), and GAPDH, in R059 parental versus two independent shRNA-mediated knockdown clones of either CBP or p300.
- **(B)** Representative Western blots of H3K27Lac (arrow), H3K27ac (arrow), and total H3 in R059 parental versus two independent shRNA-mediated knockdown clones of either CBP or p300.
- **(C)** Representative Western blots of CBP (arrow), p300 (arrow) and GAPDH in BT245 (left) and DIPG007 (right) treated with 0, 10, or 100nM of CBPD-409 for 48 hr.
- **(D)** Representative western blots of H3K27Lac (arrow), H3K27ac (arrow), and total H3 in BT245 (left) and DIPG007 (right) treated with 0, 10, or 100nM of CBPD-409 for 48 hr.
- **(E)** Representative Western blots of H3K27M (arrow) and total H3 in DIPG007 cells treated with 0, 10, or 100 nM of CBPD-409 for 48h.
- **(F)** Schema showing the cell lines and conditions (DIPG007 treated with vehicle or 100nM CBPD409 for 48 hours; BT245 treated with vehicle or 100nM CBPD409 for 48 hours) used in H3K27Lac and H3K27ac ChIP-seq and downstream analysis.
- **(G)** Box-and-whisker plots showing normalized H3K27Lac read counts (counts per million) distribution in DIPG007 vehicle versus CBPD-409 treated (top), and BT245 vehicle versus CBPD-409 treated (bottom). Data analyzed using Mann-Whitney U test. Boxes show the interquartile range (IQR), with median lines; whiskers extend to the most extreme values within 1.5 × IQR of the box, with p-values shown above each comparison. Cells were treated with 100 nM CBPD-409 for 48h.
- **(H)** Boxen plots showing the quartile distribution of H3K27Lac enrichment and medians enrichment score across LINE, DNA, LTR, and SINE classes of transposable elements in in DIPG007 vehicle versus CBPD-409 treated (top), and BT245 vehicle versus CBPD-409 treated (bottom). Central boxes span the 25th–75th percentiles, with progressively narrower boxes representing successive halvings of each tail. Statistical significance was determined using a two-sided Mann–Whitney U tests, with p-values displayed above each comparison. Cells were treated with 100 nM CBPD-409 for 48h.
- **(I)** Heatmaps showing H3K27Lac ChIP-seq enrichment centered on peak (±3 kb) at promoters and enhancers for DIPG007 and BT245 treated with vehicle or CBPD-409. Individual peaks are displayed as rows and ordered by signal intensity. Red heatmaps denote genomic regions with higher H3K27Lac occupancy, blue heatmaps denote genomic regions with lower H3K27Lac occupancy, and the corresponding color scales are displayed on the right. Metaplots above each heatmap depict the mean signal across all peaks. Numbers on the side of heatmaps indicate percent change within each comparison; peaks with similar signal are not shown (methods). Cells were treated with 100 nM CBPD-409 for 48h.
- **(J)** Schema illustrating conditions compared to assess H3K27Lac similarities in promoter and enhancer occupancy. <u>Top</u>: The comparisons in DIPG007 cells shH3-3A/parental versus CBPD-409/parental. <u>Bottom</u>: The comparisons in BT245 cells CRISPR/parental versus CBPD-409/parental treatment.
- **(K)** Heatmaps showing H3K27Lac ChIP-seq enrichment centered on peak (±3 kb) at promoters and enhancers for DIPG007 (parental, shH3-3A, and CBPD-409 treatment), and BT245 (parental, CRISPR, and CBPD-409 treatment). Individual peaks are displayed as rows and ordered by signal intensity. Red heatmaps denote genomic regions with higher H3K27Lac occupancy, blue heatmaps denote genomic regions with lower H3K27Lac occupancy, and the corresponding color scales are displayed on the right. Metaplots above each heatmap depict the mean signal across all peaks. Numbers on the side of heatmaps indicate percent change within each comparison; peaks with similar signal are not shown (methods). Cells were treated with 100 nM CBPD-409 for 48h.
- **(L)** Dot plot showing H3K27Lac-dominant significantly enriched pathways identified by gene set enrichment analysis (GSEA) in DIPG007; shH3-3A and CBPD-409 treated cells (top) and H3K27M cells (bottom). X-axis is the proportion of genes in each pathway; y-axis shows specific pathways (red highlighted pathways are common between the two cell lines); dot size indicates the number of genes in each pathway, and color intensity represents the adjusted *p*-value.
- **(M)** Dot plot showing H3K27Lac-dominant significantly enriched pathways identified by gene set enrichment analysis (GSEA) in BT245; CRISPR and CBPD-409 treated cells (top) and H3K27M cells (bottom). X-axis is the proportion of genes in each pathway; y-axis shows specific pathways (red highlighted pathways are common between the two cell lines); dot size indicates the number of genes in each pathway, and color intensity represents the adjusted *p*-value. Figures in panels F and J were generated using biorender.

**Figure S5.**
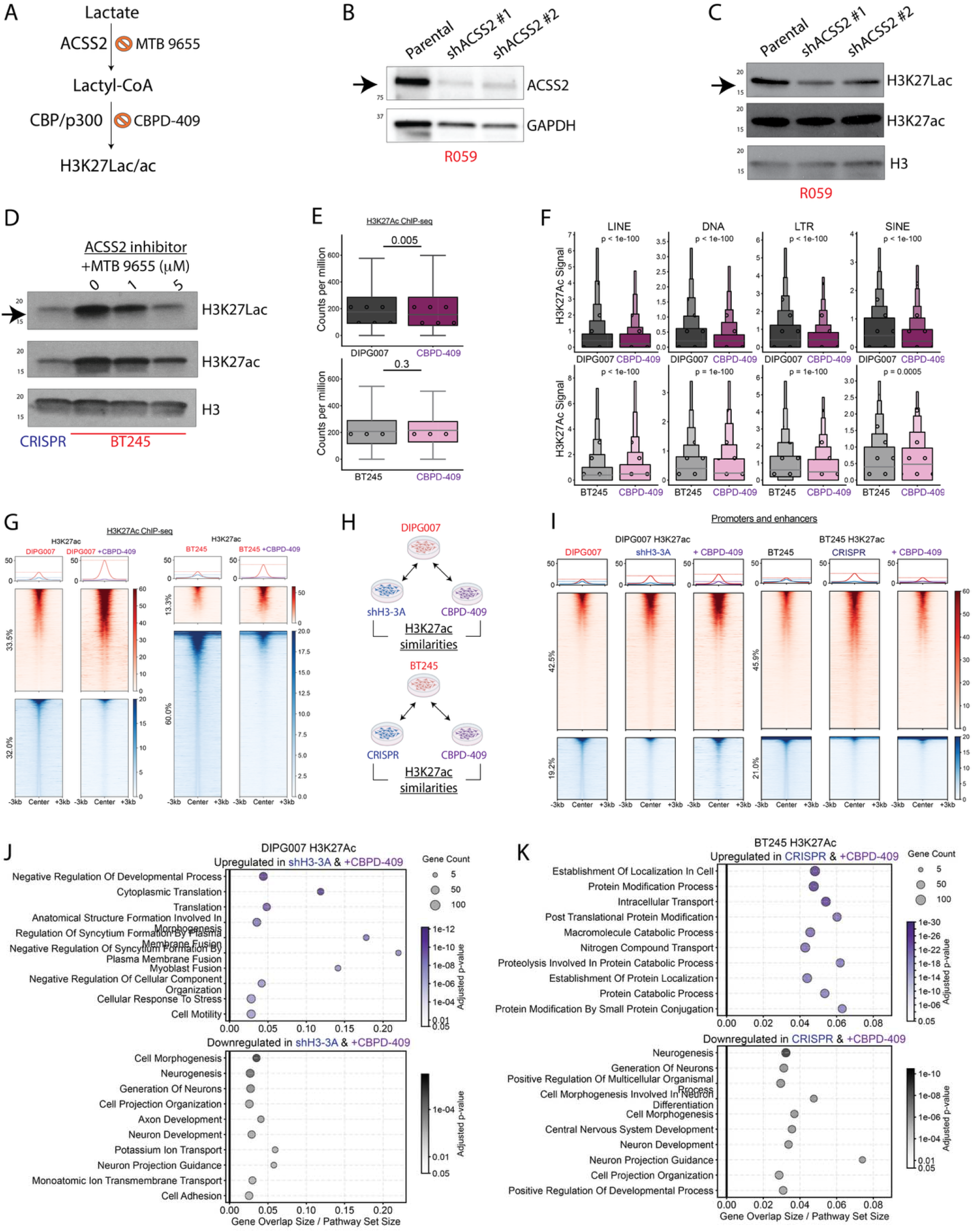
CBP/p300 inhibition simultaneously lowers H3K27Lac and H3K27ac and decreases their enrichment at ERV elements (related to figure 6) - **(A)** Schematic illustrating enzymes and inhibitors at two critical nodes critical for H3K27Lac deposition. ACSS2 converts lactate to lactyl-CoA, and CBP/p300 deposits both H3K27ac and H3K27Lac marks.
- **(B)** Representative Western blots of ACSS2 (arrow) and GAPDH in R059 parental and two independent shRNA-mediated knockdown clones of ACSS2.
- **(C)** Representative Western blots of H3K27Lac (arrow), H3K27ac, and total H3 in R059 parental and two independent shRNA-mediated knockdown clones of ACSS2.
- **(D)** Representative Western blots of H3K27Lac (arrow), H3K27ac, and total H3 in BT245 treated with 0, 1, or 5 uM of the ACSS2 inhibitor MTB9655 for 48hr. Isogenic CRISPR corrected cells used as control.
- **(E)** Box-and-whisker plots showing normalized H3K27ac read counts (counts per million) distribution in DIPG007 vehicle versus CBPD-409 treated (top), and BT245 vehicle versus CBPD-409 treated (bottom). Data analyzed using Mann-Whitney U test. Boxes show the interquartile range (IQR), with median lines; whiskers extend to the most extreme values within 1.5 × IQR of the box, with p-values shown above each comparison. Cells were treated with 100nM CBPD-409 for 48h.
- **(F)** Boxen plots showing the quartile distribution of H3K27ac enrichment and median enrichment score across LINE, DNA, LTR, and SINE classes of transposable elements in in DIPG007 vehicle versus CBPD-409 treated (top), and BT245 vehicle versus CBPD-409 treated (bottom). Central boxes span the 25th–75th percentiles, with progressively narrower boxes representing successive halving of each tail. Statistical significance was determined using a two-sided Mann– Whitney U tests, with p-values displayed above each comparison. Cells were treated with 100nM CBPD-409 for 48h.
- **(G)** Heatmaps showing H3K27ac ChIP-seq enrichment centered on peaks (±3 kb) at promoters and enhancers for DIPG007 and BT245 with or without CBPD-409 treatment. Individual peaks are displayed as rows and ordered by signal intensity. Red heatmaps denote genomic regions with higher H3K27ac occupancy, blue heatmaps denote genomic regions with lower H3K27ac occupancy, and the corresponding color scales are displayed on the right. Metaplots above each heatmap depict the mean signal across all peaks. Numbers on the side of heatmaps indicate percent change within each comparison; peaks with similar signal are not shown (methods). Cells were treated with 100nM CBPD-409 for 48h.
- **(H)** Schema illustrating conditions compared to assess H3K27ac similarities in promoter and enhancer occupancy. <u>Top</u>: The comparisons in DIPG007 cells shH3-3A/parental versus CBPD-409/parental. <u>Bottom</u>: The comparisons in BT245 cells CRISPR/parental versus CBPD-409/parental treatment.
- **(I)** Heatmaps showing H3K27ac ChIP-seq enrichment centered on peak (±3 kb) at promoters and enhancers for DIPG007 (parental, shH3-3A, and CBPD-409 treatment), and BT245 (parental, CRISPR, and CBPD-409 treatment). Individual peaks are displayed as rows and ordered by signal intensity. Red heatmaps denote genomic regions with higher H3K27ac occupancy, blue heatmaps denote genomic regions with lower H3K27ac occupancy, and the corresponding color scales are displayed on the right. Metaplots above each heatmap depict the mean signal across all peaks. Numbers on the side of heatmaps indicate percent change within each comparison; peaks with similar signal are not shown (methods). Cells were treated with 100 nM CBPD-409 for 48h.
- **(J)** Dot plot showing H3K27Ac-dominant significantly enriched pathways identified by gene set enrichment analysis (GSEA) in DIPG007; shH3-3A and CBPD-409 treated cells (top) and H3K27M cells (bottom). X-axis is the proportion of genes in each pathway; y-axis shows specific pathways (red highlighted pathways are common between the two cell lines); dot size indicates the number of genes in each pathway, and color intensity represents the adjusted *p*-value.
- **(K)** Dot plot showing H3K27Ac-dominant significantly enriched pathways identified by gene set enrichment analysis (GSEA) in BT245; CRISPR and CBPD-409 treated cells (top) and H3K27M cells (bottom). X-axis is the proportion of genes in each pathway; y-axis shows specific pathways (red highlighted pathways are common between the two cell lines); dot size indicates the number of genes in each pathway, and color intensity represents the adjusted *p*-value. Figures in panel H were generated using biorender.

**Figure S6.**
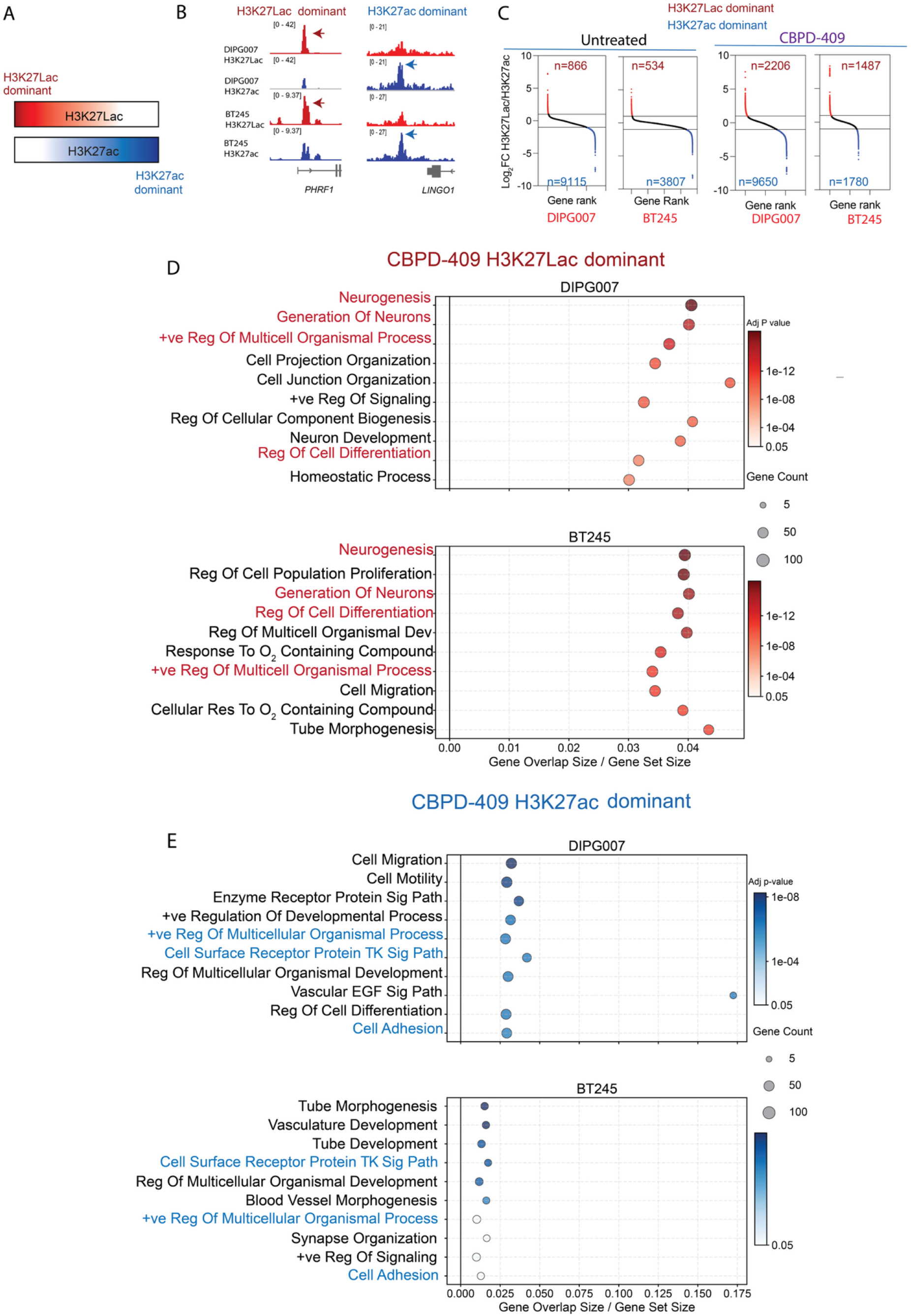
CBP/p300 inhibition simultaneously lowers H3K27Lac and H3K27ac and decreases their enrichment at ERV elements (related to figure 6) - **(A)** Schema showing comparative H3K27Lac versus H3K27ac dominant genomic regions. Color gradients represent genomic distribution of ChIP-seq signal intensity for H3K27Lac and H3K27ac. Lighter shading corresponds to lower signals, and darker shading corresponds to higher signals.
- **(B)** <u>Left</u>: Representative ChIP tracks showing H3K27Lac dominant *PHRF1* genomic locus. H3K27Lac (red) and H3K27ac (blue) in DIPG007 (top two tracks) and BT245 (bottom two tracks). <u>Right</u>: Representative ChIP tracks showing H3K27ac dominant *LINGO1* genomic locus. H3K27Lac (red) and H3K27ac (blue) in DIPG007 (top two tracks) and BT245 (bottom two tracks).
- **(C)** Ranked distribution of genomic loci according to the log2 fold change of H3K27Lac/H3K27ac in DIPG007, BT245, DIPG007 treated with 100nM CBPD409, and BT245 treated with 100nM CBPD409 (left to right). Values greater than 1 represent genomic loci with H3K27Lac dominance (red). Values less than −1 represent genomic loci with H3K27ac dominance (blue). Black dots represent genomic regions with intermediate scores contained within dominance threshold (horizontal black lines).
- **(D)** Dot plot showing H3K27Lac-dominant significantly enriched pathways identified by gene set enrichment analysis (GSEA) in DIPG007 (top) and BT245 (bottom). X-axis is the proportion of genes in each pathway; y-axis shows specific pathways (red highlighted pathways are common between the two cell lines); dot size indicates the number of genes in each pathway, and color intensity represents the adjusted *p*-value.
- **(E)** Dot plot showing H3K27ac-dominant significantly enriched pathways identified by GSEA in DIPG007 (top) and BT245 (bottom). X-axis is the proportion of genes in each pathway; y-axis shows specific pathways (blue highlighted pathways are common between the two cell lines); dot size indicates the number of genes in each pathway, and color intensity represents the adjusted *p*-value.

We assessed genomic distribution of H3K27Lac and H3K27ac in H3K27M cells treated with or without CBPD-409 using chromatin immunoprecipitation sequencing (ChIP-seq) (**Fig 6F**). We used ChIP-grade antibodies specifically designed to differentiate H3K27Lac from H3K27ac genomic distribution.^56–59^ CBPD-409 reduced global genomic H3K27Lac (**Fig 6G**). As observed with suppression of mutant H3K27M (**Fig 2**), CBPD-409 lowered H3K27Lac at intergenic regions including LINE, SINE, LTR, and DNA classes of ERV transposable repetitive elements in DMG cells (**Fig 6H**). Additionally, CBPD-409 treatment was associated with substantial redistribution of H3K27Lac at promoters and enhancers (**Fig 6I**). We noted similar results with H3K27ac where CBPD-409 also reduced global H3K27ac and at intergenic regions, and remodeled H3K27ac at promoters and enhancers (**Fig S5E-G**). We overlapped changes in genomic H3K27Lac in shH3-3A versus DIPG007, and CBPD-409 versus DIPG007 vehicle treated cells; and CRISPR versus BT245, and CBPD-409 versus BT245 vehicle treated cells (**Fig 6J**). These studies showed similar changes in H3K27Lac distribution with either H3K27M suppression or CBPD-409 treatment in both cell lines (**Fig 6K**). Similar results were noted with H3K27ac distribution in these conditions (**Fig S5H-I**).

We next compared genomic enrichment of H3K27Lac with H3K27ac at promoters and enhancers in both cell lines treated with or without CBPD-409. Because histone H3 is composed of greater than 16 alleles, global assessment of post translational modifications at a given H3K residue reflects the combined state of multiple H3 gene-encoded modified lysine residues. For example, hepatic stellate cells exhibit overlap in genomic enrichment of both H3K18Lac with H3K18ac.^43^ However, several genomic loci show greater H3K18Lac versus H3K18ac enrichment, and *vice versa*.^60^ We determined if similar biologic principles apply to H3K27Lac and H3K27ac in H3K27M cells (**Fig S6A**). While the two marks showed overlap, we identified gene loci with H3K27Lac high, but low H3K27ac (H3K27Lac dominant, >1 Log_2_ fold change H3K27Lac/H3K27ac) and *vice versa* (H3K27ac dominant, < −1 Log_2_ fold change H3K27Lac/H3K27ac (**Fig S6B-C**). We performed GSEA with H3K27Lac-dominant and H3K27ac-dominant regions in CBPD-409 versus vehicle treated cells. We noted that H3K27Lac-dominant regions corresponded mainly to neuro-developmental and regulation of differentiation-related pathways in both DIPG007 and BT245 H3K27M cells (**Fig S6D**). H3K27ac-dominant regions corresponded to cell adhesion and receptor tyrosine kinase pathways in both cell lines (**Fig. S6E**).

Overall, CBPD-409 treatment resulted in simultaneously lowering of global and intergenic H3K27Lac and H3K27ac and redistribution of both marks at promoters and enhancers in H3K27M cells related to differentiation and other pathways. Moreover, CBPD-409 treatment partially phenocopied changes in H3K27Lac and H3K27ac genomic distribution on H3K27M suppression. We next determined function significance of CBPD-409 treatment in both cell lines and animal models.

### CBPD-409 suppresses MES- and OPC 2-like signatures, upregulates PTEN, and is therapeutic in preclinical H3K27M models

We assessed the relationship between changes in genomic H3K27Lac distribution with gene expression to determine functional relevance of CBPD-409 treatment. Regions with lower H3K27Lac enrichment and decreased gene expression corresponded to pathways related to EMT, hypoxia, and neurodevelopmental genes (**Fig 7A**). Amongst the downregulated genes, key neuronal developmental factors (*OLIG1, OLIG2,* and *CRB2*); EMT genes (*NNMT, SNTB1, COL5A1, ECM1, PDGFRB, ECSCR,* and *ELN*) and hypoxia signature genes (*HK2*, *ANXA2*, and *ATF4*3) showed both decreased H3K27Lac enrichment and gene expression (**Fig 7B-C, S7A**). Protein levels of several of these factors including OLIG2, HK2, and OLIG1 were reduced in H3K27M patient-derived cells treated with CBPD-409 versus vehicle (**Fig 7D and Fig S7B**). H3K27ac also showed reduced enrichment at these genes but was more variable (**Fig 7B, S7C**). In contrast, upregulated genes that were enriched for H3K27Lac corresponded to several tumor suppressor genes, including *PTEN, RB1, ZBTB20, NF1*, and *BCL10* (**Fig 7E-G**). PTEN protein levels were increased on treatment with CBPD-409 in two independent H3K27M cell lines (**Fig 7H**). H3K27ac enrichment was variably increased at these gene loci (**Fig 7F and Fig S7D**).

**Figure 7.**
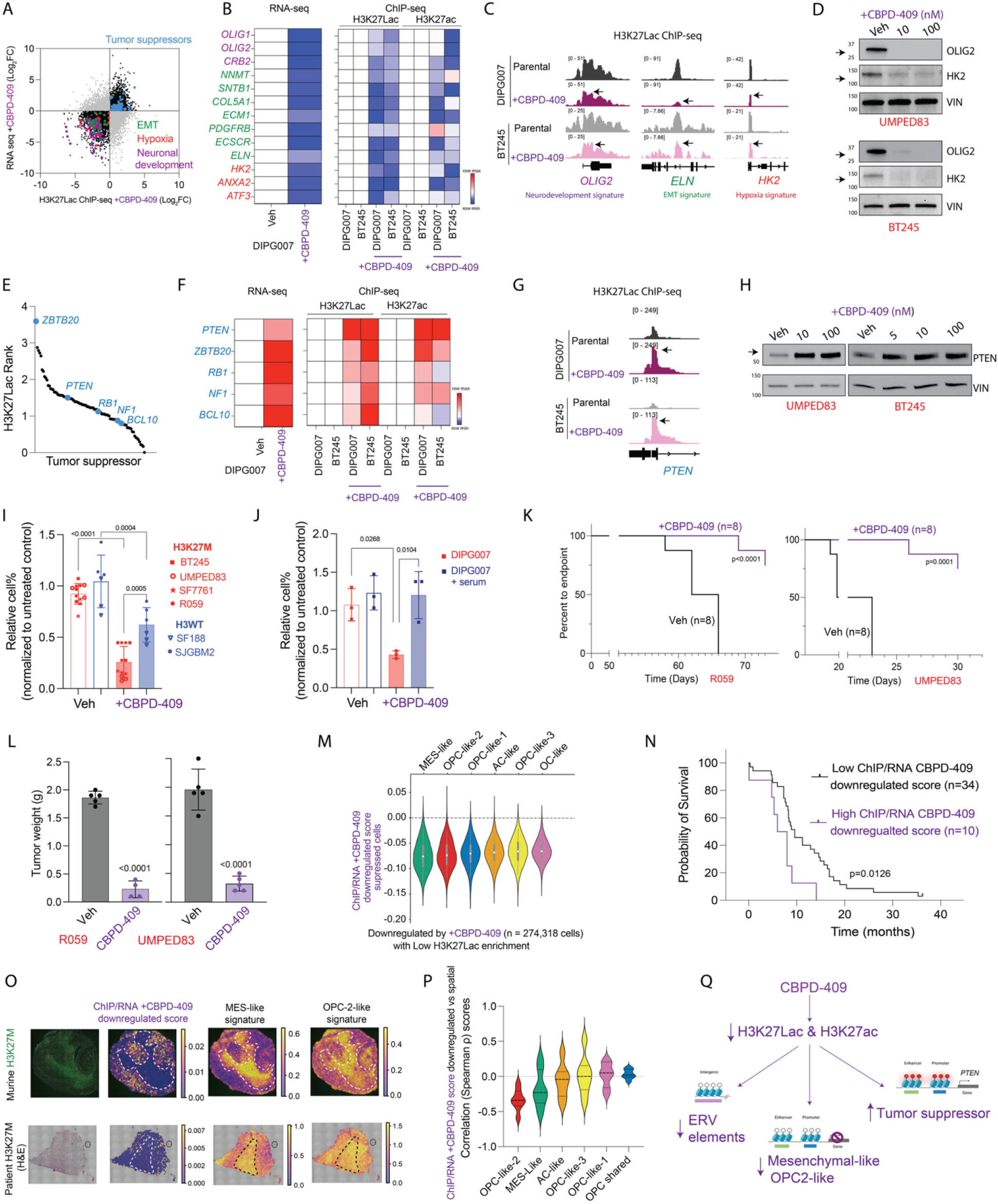
CBPD-409 suppresses MES- and OPC 2-like signatures, upregulates PTEN, and is therapeutic in preclinical H3K27M models. - **(A)** Scatter plot showing H3K27Lac ChIP-seq occupancy (Log_2_FC; x-axis) and differential gene expression (Log_2_FC; y-axis) in DIPG007+CBPD409 relative to vehicle treated cells. Genes in the upper right quadrant exhibit increases in H3K27Lac occupancy and transcript expression. Genes in the lower left quadrant exhibit decreases in H3K27Lac occupancy and transcript expression. Genes mapping to “tumor suppressors (blue),” “EMT (green),” “hypoxia (red),” and “neuronal development (purple)” pathways are indicated (GSEA).
- **(B)** <u>Left</u>: heatmap showing normalized expression levels (RNA-seq) of genes in DIPG007 treated with vehicle or CBPD-409 (100nM, 48h). <u>Right</u>: Heatmap showing H3K27Lac and H3K27ac enrichment at indicated gene loci in DIPG007, BT245, DIPG007+CBPD409, and BT245+CBPD409 conditions. Genes mapping to “EMT (green),” “hypoxia (hypoxia),” and “neuronal development (purple)” are indicated. The color scale reflecting the relative signal enrichment is displayed on the right.
- **(C)** Representative showing H3K27Lac enrichment at the *OLIG2* (neuro-developmental signature gene), *HK2* (hypoxia signature gene) and *ELN* (EMT signature gene) gene loci in DIPG007, DIPG007+CBPD, BT245, and BT245+CBPD conditions.
- **(D)** Representative Western blots of OLIG2 and HK2 in UMPED83 (top) and BT245 (bottom) treated with vehicle, 10nM, or 100nM of CBPD-409.
- **(E)** Ranked tumor suppressor by H3K27Lac genomic enrichment in CBPD-409 (100nM, 48h) versus vehicle treated DIPG007 cells. Each point denotes a tumor suppressor gene (x-axis), with y-axis indicating the log_2_FC in genomic H3K27Lac ChIP-seq signal at the associated gene locus.
- **(F)** <u>Left</u>: heatmap showing normalized expression levels (RNA-seq) of representative tumor suppressor genes in DIPG007 treated with vehicle or CBPD-409 (100nM, 48h). <u>Right</u>: Heatmap showing H3K27Lac and H3K27ac enrichment at indicated gene loci in DIPG007, BT245, DIPG007+CBPD-409, and BT245+CBPD-409 conditions. The color scale reflecting the relative signal enrichment is displayed on the right.
- **(G)** Representative H3K27Lac ChIP tracks at the *PTEN* (left) tumor suppressor locus in DIPG007 versus DIPG007+CBPD (top), and BT245 versus BT245+CBPD (bottom) conditions.
- **(H)** Representative Western blots of PTEN (arrow) and GAPDH in UMPED83 (left) and BT245 (right) treated with vehicle or indicated concentrations of CBPD-409 for 48h.
- **(I)** Relative cell number (normalized to untreated control, y-axis) in vehicle or CBPD-409 treated (100nM, 48h) H3K27M cells (BT245, UMPED83, SF7761, and R059) and H3WT cells (SF188 and SJGBM2). Data are shown as mean ± SD and analyzed with one-way ANOVA with Tukey’s multiple comparisons using 95% C.I., n=3 for each cell line, P values are indicated.
- **(J)** Relative cell number (normalized to untreated control, y-axis) in vehicle or CBPD-409 treated DIPG007 and serum differentiated DIPG007 (cultured in 10% FBS for 2 weeks). Data shown as mean ± SD and analyzed by using two-way ANOVA with Šídák’s multiple comparisons using 95% C.I., n=3 for each cell line, P values are indicated.
- **(K)** Kaplan-Meier plots showing the percentage of mice remaining below the study endpoint (4x increase in tumor volume, y-axis) over time (x-axis) in R059 (left) and UMPED83 (right) H3K27M in vivo models. Mice were randomly assigned to vehicle (black) or CBPD-409 treatment (purple) groups after tumor development. Mice were treated with vehicle or 3mg/kg CBPD-409 by oral gavage for 2 weeks, 3 days per week. Statistical analysis was performed using Log-rank test with 95% C.I. P values are indicated.
- **(L)** Bar graphs showing tumor weights harvested at the study endpoint from UMPED83 (left) and R059 (right) treated with CBPD-409 (purple, n=4) or vehicle (black, n=5) from G. Data shown as mean ± SD and analyzed by unpaired, two-tailed, two-sided t-test with 95% C.I.
- **(M)** Violin plots showing the projected distribution of cells (wishbone plot in fig 4F) with genes that showed decreased H3K27Lac enrichment and downregulated expression in DIPG007 CBPD treated cells. The x-axis shows the indicated cell states. The width of each violin represents the density of observations at each value. The white dot denotes the median, and the gray box indicates the interquartile range (IQR), with whiskers representing the distribution range. The dashed horizontal line indicates 0, corresponding to no change in gene regulation.
- **(N)** Genes that showed decreased H3K27Lac enrichment and downregulated expression in DIPG007 CBPD-409 treated cells were derived from published H3K27M clinical data sets (Macaky et al. *Cancer Cell* 2017). Gene expression data were stratified into two H3K27M patient groups by unbiased K-means clustering into high (n=10, purple) versus low (n=36, black) groups. Kaplan-Meier survival analysis (with 95% C.I.) of published patient overall survival was performed between the two groups. P value is indicated.
- **(O)** Representative spatial maps from tissue samples (top row: mouse H3K27M tumors; bottom row patient H3K27M sample, data derived from Colot et al. 2026).
  – The first column shows H3K27M stained murine H3K27M tumor tissue or hematoxylin and eosin (H&E) stained patient H3K27M DMG.
  – The second column shows spatial enrichment of CBPD ChIP/RNA score (genes with downregulated genomic H3K27Lac and expression).
  – The third and fourth columns show spatial enrichment and MES-like and OPC 2-like gene signatures. Color intensity represents the magnitude of the corresponding molecular score, with scales shown next to each panel.
- **(P)** Violin plots showing correlation of integrated ChIP/RNA Spearman ρ score with spatial transcriptomics of indicated cell states (x-axis) after 100nM CBPD-409 treatment. The width of each violin represents the density of observations at each value. The black dashed horizontal lines within each violin indicate the 25th, 50th (median), and 75th percentiles, respectively. The height of each violin indicates the range of distribution.
- **(Q)** Scheme showing CBPD-409 lowering both genomic H3K27Lac and H3K27ac at ERV elements; and OPC2-like, and MES-like gene signature genes. This was accompanied by H3K27Lac activation of tumor suppressors including PTEN. Figures in panel Q were generated using biorender.

**Figure S7.**
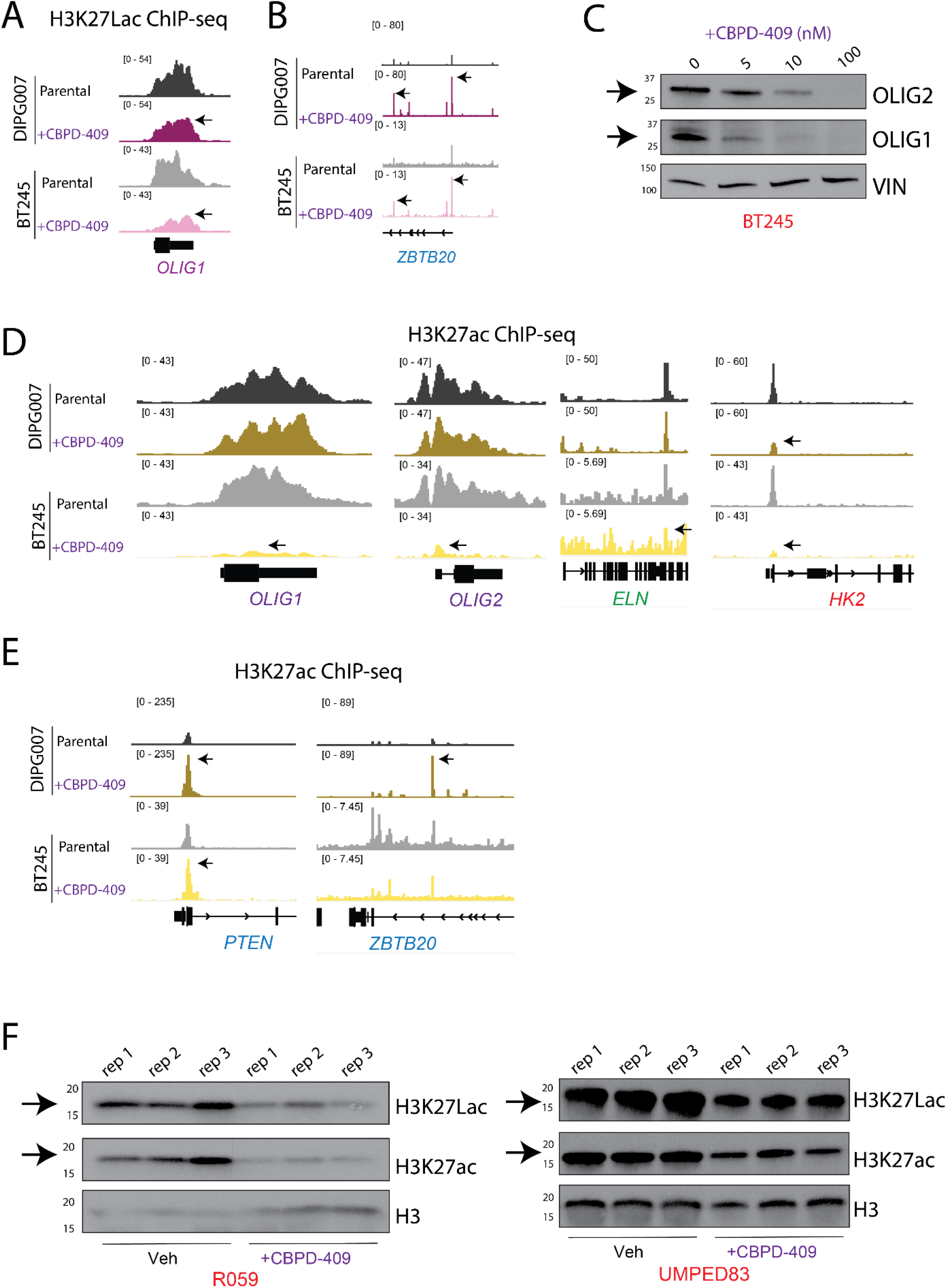
CBPD-409 suppresses MES- and OPC 2-like signatures, upregulates PTEN, and is therapeutic in preclinical H3K27M models (related to Figure 7) - **(A)** Representative H3K27Lac ChIP tracks at the *OLIG1* locus in DIPG007, DIPG007+CBPD, BT245, and BT245+CBPD conditions.
- **(B)** Representative H3K27Lac ChIP tracks at the *ZBTB20* tumor suppressor locus in DIPG007 versus DIPG007+CBPD (top), and BT245 versus BT245+CBPD (bottom) conditions.
- **(C)** Representative Western blots of OLIG2, OLIG1 and vinculin in BT245 treated with vehicle, 5nM, 10nM, or 100nM of CBPD-409 conditions for 48h.
- **(D)** Representative H3K27ac ChIP tracks at the *OLIG1*, *OLIG2*, *ELN*, and *HK2* loci in DIPG007, DIPG007+CBPD, BT245, and BT245+CBPD conditions.
- **(E)** Representative H3K27ac ChIP tracks at the *PTEN* (left) and *ZBTB20* (right) loci in DIPG007 versus DIPG007+CBPD (top two tracks), BT245 versus BT245+CBPD (bottom two tracks).
- **(F)** Western blots showing H3K27Lac (arrow), H3K27ac (arrow), and total H3 in R059 (left) and UMPED83 (right) harvested tumor tissues treated with vehicle or CBPD-409 from *in vivo* experiments (n=3, each).

We tested the therapeutic efficacy of CBPD-409 *in vitro.* Toxicity studies showed that while both H3K27M and H3WT patient-derived cells were sensitive to CBPD-409 treatment, H3K27M versus H3WT cells showed greater sensitivity (**Fig 7I**). Moreover, H3K27M cells cultured in the presence of serum, which showed lower H3K27Lac (**Fig 4J**), lost CBPD-409 sensitivity (**Fig 7J**). Pharmacokinetic studies demonstrated minimal brain exposure to CBPD-409, with brain concentrations below the limit of quantification (<5 ng/g) at 7 hours following IV or PO administration despite measurable plasma concentrations. Therefore, we tested proof-of-principle efficacy in two independent H3K27M flank xenograft models. Therefore, we tested proof-of-principle efficacy of CBPD-409 in two independent H3K27M in vivo flank animal models. CBPD-409 reduced tumor burden in both models (**Fig 7K-L**). Moreover, H3K27Lac and H3K27ac showed a global reduction in vehicle versus CBPD-409 treated tumor samples (**Fig S7E**).

We assessed the relationship between H3K27M tumor cell states and CBPD-409 downregulated genes (showing both lower H3K27Lac enrichment and downregulated by CBPD-409 versus vehicle treated cells). CBPD-409 suppressed genes showed greatest reduction in OPC-like-2 and MES-like cell states (**Fig 7M**). We also assessed the potential clinical significance of these CBPD-409 suppressed genes. We used K-means clustering of CBD-409 suppressed genes to stratify H3K27M DMG (n=46) patients into high (n=10) versus low (n=34) expression groups.^48^ High versus low expression of CBPD-409 suppressed genes showed a significant (p=0.0126) worse overall survival outcome in H3K27M DMGs (**Fig 7N**). We confirmed our cell-state findings using published spatial transcriptomic data.^28^ We noted the largest negative correlation between spatial distribution of CBPD-409 suppressed genes with MES-like and OPC-like-2 cell state signatures (**Fig 7O-P**). In summary, CBPD-409 treatment: (a) globally lowered H3K27Lac and H3K27ac, (b) decreased H3K27Lac and H3K27ac enrichment at intergenic regions including ERV elements, (c) upregulated expression of tumor suppressors like PTEN, and (d) downregulated key neuronal stem cell/ OPC-like and MES-like signatures. (**Fig 7Q**).

## Discussion

Growing evidence supports metabolic coupling with epigenetically driven cell differentiation, whereby changes in nutrient availability and their utilization can directly influence gene transcription and cell fate decisions.^61^ In this study we determined if the metabolite lactate epigenetically regulates tumor cell state heterogeneity in H3K27M DMGs. We discovered that H3K27Lac was the only histone lysine-lactyl mark, amongst the ten we assessed, that was increased in H3K27M versus H3WT or H3G34-mutant cells. Using patient MRS imagining, cell line and animal models, and tumor spatial transcriptomics, we demonstrate that: (a) lactate levels are heterogeneous within H3K27M DMG; (b) exogenous lactate increases global H3K27Lac levels, corresponding to activation of key OPC-related and MES signature genes; and (c) lactate driven gene expression relates with poor overall outcome in patient tumors. Because of shared enzymatic machinery, targeting CBP/p300 presented a dual-pronged strategy to lower both H3K27Lac and H3K27ac. This resulted in greatest suppression of MES- and OPC-like-2 cell states, and upregulation of the tumor suppressor PTEN.

A heterogeneous population of cells, reflecting a spectrum of cell differentiation states, is found in the H3K27M DMG tumor microenvironments. These include OC-like, OPC-like-1, OPC-like-2, OPC-like-3, and AC-like cells. OPC-like-2 and OPC-like-3 versus OPC-like-1 cells correspond to more immature malignant oligodendrocyte precursors.^24^ Aggressive MES-like H3K27M cells are linked to EMT states, hypoxia, invasion, inflammatory signaling, and extracellular-matrix remodeling.^23,24,28,29,62–64^ Our data suggest that exogenous lactate can epigenetically enhance key OPC and MES related signature genes. Amongst OPC-like signatures, overlap of lactate driven genes enriched by H3K27Lac with single cell analyses from patient tumors corresponded mainly to OPC-like-2 subsets. Interesting, lactate plays a role in oligodendrocyte maturation and remyelination.^65–67^ For example, ablating LDHA in oligodendrocytes resulted in defective oligodendrocyte development leading to dysmyelination.^65^ In line with these results, we noted that normal OPC versus AC cells isolated from wildtype mice showed higher H3K27Lac levels. Together these studies and our data suggest that OPC-like H3K27M tumor cells may hijack developmentally conserved programs, including H3K27-lactylation to promote their aggressive nature.

While lactate driven EMT is observed in several cancers, epigenetic mechanisms that regulate this phenomenon are poorly understood.^68–71^ We identified TGFβ1 as a downstream target of H3K27Lac in response to exogenous lactate and hypoxia. TGFβ1 is recognized as a master EMT regulator with known roles in activating a coordinated cell-state program that remodels the extracellular matrix.^72^ This finding provides a potential mechanism for a H3K27Lac-mediated and TGFβ1-driven activation of a mesenchymal transcriptional programs that contribute to aggressive DMG biology.

Previous studies have demonstrated that although global levels of H3K27ac are high, active enhancers containing H3K27M compared to H3WT exhibit lower H3K27ac enrichment in human hindbrain cell models.^11, 21^ The opposite pattern is observed at intergenic regions with greater H3K27ac pervasive deposition at endogenous retroviral (ERV) and other intergenic areas in H3K27M versus shH3-3A cells.^21^ This widespread intergenic deposition of H3K27ac has been proposed to derepress repetitive elements and activate innate immunity through “viral mimicry” in DMG cells.^13^ Our H3K27Lac results are similar to these studies where suppression of H3K27M by knockdown or CRISPR correction led to a decrease in H3K27Lac at ERV LINE, SINE and LTR classes of ERV elements. No consistent changes were observed at ERV regions on exogenous lactate treatment. However, CBPD-409, similar to H3K27M suppression, reduced H3K27Lac at these ERV elements warranting future work on the functional significance of this finding.

Histone lysine residues can be modified by a variety of acylation groups including lactylation and acetylation.^39,44,73,74^ Multiple acylation marks globally coexist on lysine residues of histone H3 because it is composed of more than 16 alleles, reflecting an overall combined state.^39,44,73,74^ For example, H3K18, H3K23, H3K27 can each bear multiple acylation marks including acetylation, lactylation and propionylation. Moreover, these marks can show overlap in genomic distribution.^60,75,76^ This is reflected globally (as assessed by Western blotting or ChIP-seq/ CUT&RUN/ CUT&Tag) by multiple modifications at any given H3K residue. For example, H3K18 exhibits both lactyl (H3K18Lac) and acetyl (H3K18ac) modifications in hepatocyte stellate cells.^39,60^ By way of illustration, H3K18Lac and H3K18ac co-occupy genomic regions in hepatic stellate cells, but bear genes with comparatively dominant H3K18Lac or H3K18ac enrichment. We noted similar patterns of H3K27Lac and H3K27ac enrichment with H3K27Lac dominant genes corresponding mainly to neurodevelopmental pathways.

We took advantage of similar enzymatic machinery that mediates histone acetylation and lactylation for potential therapeutic development. Targeting CBP/p300 using CBPD-409 simultaneously lowered global H3K27Lac and H3K27ac levels. CBPD-409 lowered genomic H3K27Lac and expression of key genes related to EMT, OPC and MES-like tumor cell states. CBP/p300 degradation also activated tumor suppressors including PTEN. This bidirectional effect of suppressing key neuronal developmental and tumor cell states, while concomitantly upregulating tumor suppressors is unexpected. Our discoveries suggest that inhibiting CBP/p300 leads to remodeling of the H3K27M regulatory landscape and may warrant further investigation.

Our study has several limitations that need to be considered. Lactate can decorate several histone lysine residues. We chose to focus on H3K27Lac as it was the only lactyl-mark that was increased in H3K27M versus H3WT or H3G34-mutant cells.

However, we cannot rule out the effects of exogenous lactate treatment on other histone lysine residues including H3K18. Moreover, non-histone protein lactylation also plays an important role in cancer. For example, lactylation of the nucleoside diphosphate kinase NME1 increases the synthesis of nucleoside triphosphates in H3K27M DMG cells.^40^ Similarly, inhibition of CBP/p300 can simultaneously lower multiple histone acyl modifications. While our data suggests that CBPD-409 reduces both H3K27Lac and H3K27ac, we cannot rule out suppression of other histone lysine acyl modifications. CBPD-409 did not penetrate the brain leading to proof-of-principle tumor flank *in vivo* experiments. Nevertheless, our data suggest that H3K27Lac plays an important role in H3K27M DMGs. Furthermore, reduction of multiple histone lysine acyl modifications by CBPD-409 can be viewed as an advantage that tackles the tumor on multiple levels requiring further investigation into enhancing brain penetration and dissecting out mechanisms of action.

In summary, this study highlights lactate as a crucial factor in epigenetically regulating tumor cell state heterogeneity in H3K27M DMGs via H3K27Lac. providing a potential mechanism by which the Warburg effect is coupled to cell state tumor heterogeneity in H3K27M DMGs. We found elevated H3K27Lac levels in H3K27M cells, and exogenous lactate increased H3K27Lac activation of key genes linked to OPC and MES-like cell fates correlating with poorer patient outcomes. Additionally, the PROTAC compound CBPD-409 reduced both H3K27Lac and H3K27ac levels and aggressive MES-like and OPC-like-2-gene signatures. Overall, our findings suggest targeting lactate-driven epigenetic changes may offer promising therapeutic avenues for H3K27M DMGs.

## Acknowledgements

This work was funded by the Sontag Foundation (S.V.), Clinical Scientist Development Award–Doris Duke Charitable Foundation (S.V.), the Hyundai Hope On Wheels Foundation (S.V.), NINDS R01NS110572 (S.V.), NCI R01CA261926 and the Julie Taubman Reys emerging scholar award from the University of Michigan Taubman Institute (S.V.). The Venneti Laboratory is/was supported by grants from Mathew Larson (S.V.), Sidney Kimmel (S.V.), St Baldrick’s (S.V.), Claire McKenna (S.V.), Chad Tough (S.V.), Alex’s Lemonade Stand (S.V.), Storm The Heavens (S.V.) and the University of Michigan Chad Carr Pediatric Brain Tumor Initiative (S.V.). J.L was supported by the NIH Training Program in Translational Research T32 and is currently funded by the ChadTough Defeat DIPG Foundation. We thank Drs. Cristián Valls and John Prensner for bioinformatic advice and, Dr. Nada Jabado for providing isogenic BT245 cell lines.

## Author contributions

JL and SV conceived the study, planned experiments and wrote the paper; JL, NN, SS, WT, AD, SRS, MM, KB, MW, and JB performed experiments and analyzed data; BT and SB performed and analyzed MRS data; BTM provided NCI006; ETH helped with *in vivo* CBPD-409 experiments; FM, AA, HC, MS-C, SE, FS, XC, AP, AMC, helped with H3K27Lac and H3K27ac ChIP-seq experiments; EP, DRW performed and analyzed *in vivo* tracing studies; SJ, CK helped with single cell analysis; VY provided PPK and PPW cells; RP, OBL, and KP performed and analyzed *in vitro* tracing experiments; SW and AMC provided CBPD-409.

## Methods

### Mice and housing conditions

Animal experiments were performed after approval from the University of Michigan Committee on Use and Care of Animals and were conducted as per NIH guidelines for animal welfare. All animal procedures were approved by Institutional Animal Care & Use Committee (IACUC) at the University of Michigan. Animals were housed and cared for according to standard guidelines with free access to food and water. All experiments were performed on NOD-SCID-IL2R gamma chain-deficient (NSG) mice (8–10 weeks old) with male and female animals used equally. Pregnant CD-1 International Genetic Standardization (IGS) mice and E18 embryonic mice were used for primary OPCs and ACs isolation. Animals including littermates of the same sex were randomly assigned to control or treatment conditions.

### Cell culture

All cell lines were cultured in a humidified incubator under normoxia at 37 °C with 5% CO2. For hypoxia culturing, cell lines were cultured at 37 °C with 1% oxygen and 5% CO2. Cell lines were validated by STR analysis and were routinely screened and tested negative for mycoplasma.

#### Mouse cell lines

Immortalized mNSCs (mouse neuronal stem cells) were obtained from Richard J. Gilbertson, St Jude’s Hospital. H3.3 K27M PPK, H3.1 K27M PPK, H3.3 WT PPW (PPK/PPW: PdgfrA-overexpressed and shp53 with H3.3K27M or H3.1K27M or H3WT conditions), were obtained from Vivekanand Yadav, Children’s Mercy.

H3.3K27M mNSCs, H3.1 K27M mNSCs, H3WT mNSCs, and PPK/PPW cells were cultured in Neurobasal-A (Thermo Fisher Scientific, #10888022) media supplemented with N-2 Supplement (100×; Thermo Fisher Scientific, #17502048), B-27 Supplement (50×), serum-free (Thermo Fisher Scientific, #17504044), L-glutamine (200 mmol/L; Thermo Fisher Scientific, #A2916801), 20 ng/μL human EGF (Shenandoah, #100-26), 20 ng/μL human FGF-BASIC-154aa (Shenandoah, #100-146), 2 μg/mL Heparin Solution, 0.2% (STEMCELL Technologies, Inc. #07980), 7.5% Albumin, Bovine Fraction V (mNSCs only; Dot Scientific, Inc., #DSA30075), and penicillin–streptomycin (10,000 U/mL; Thermo Fisher Scientific, #115140122).

#### Patient-derived cell lines

HSJD-DIPG007 (referred to as DIPG007) and SF7761 cells were obtained from Rintaro Hashizume, Northwestern University. BT245, BT245 CRISPR, KNS42, KNS42 CRISPR, HSJD-GBM-002, HSJD-GBM-002 CRISPR cells were obtained from Nada Jabado, University of Montreal. SU-DIPGXIII*p (referred to as DIPG13), SU-DIPG-06, SU-DIPG-13, SU-DIPG-21, SU-DIPG-25, SU-DIPG-36, SU-DIPG-38, SU-DIPG-50 cells were obtained from Michelle Monje, Stanford University. UMPED83, SJGBM2, and QCTB-R059 (referred to as R059) cells were obtained from Carl Koschmann, University of Michigan. SF188, TS543, and TS603 cells were obtained from Craig B. Thompson, Memorial Sloan Kettering Cancer Center.

DIPG007, DIPG007 shH3-3A, DIPG13, SU-DIPG-06, SU-DIPG-13, SU-DIPG-21, SU-DIPG-25, SU-DIPG-36, SU-DIPG-38, SU-DIPG-50, BT245, BT245 CRISPR, UMPED83, R059, TS603, and TS543 cells were cultured in Neurobasal-A (Thermo Fisher Scientific #10888022) and DMEM/F-12 (Thermo Fisher Scientific #11330032) media supplemented with HEPES (1 M; Thermo Fisher Scientific #15630080), sodium pyruvate (100 mmol/L; Thermo Fisher Scientific, #11360070), MEM Nonessential Amino Acids Solution (100×; Thermo Fisher Scientific, #11140050), GlutaMAX Supplement (Thermo Fisher Scientific #35050061), B-27 Supplement (50×), minus vitamin A (Thermo Fisher Scientific #12587010), 20 ng/μL human EGF (Shenandoah, #100-26), 20 ng/μL human FGF-BASIC-154aa (Shenandoah, #100-146), 10 ng/μL human PDGF-AA (Shenandoah, #100-16), 10 ng/μL human PDGF-BB (Shenandoah, #100-18), 2 μg/mL Heparin Solution, 0.2% (STEMCELL Technologies, Inc., #07980), and antibiotic-antimycotic (100×; Thermo Fisher Scientific #15240096).

SF7761 were cultured in Neurobasal-A (Thermo Fisher Scientific, #10888022) media supplemented with N-2 Supplement (100×; Thermo Fisher Scientific, #17502048), B-27 Supplement (50×), serum-free (Thermo Fisher Scientific, #17504044), L-glutamine (200 mmol/L; Thermo Fisher Scientific, #A2916801), 20 ng/μL human EGF (Shenandoah, #100-26), 20 ng/μL human FGF-BASIC-154aa (Shenandoah, #100-146), 2 μg/mL Heparin Solution, 0.2% (STEMCELL Technologies, Inc. #07980), 7.5% Albumin, Bovine Fraction V (Dot Scientific, Inc., #DSA30075), and penicillin–streptomycin (10,000 U/mL; Thermo Fisher Scientific, #115140122).

KNS42, KNS42 CRISPR, and SF188 were cultured in DMEM/F-12 (Thermo Fisher Scientific #11330032) media supplemented with 10% Fetal Bovine Serum (FBS) (Cat#VWR #89510-186), GlutaMAX Supplement (Thermo Fisher Scientific #35050061), and penicillin–streptomycin (10,000 U/mL; Thermo Fisher Scientific, #115140122).

HSJD-GBM-002 and HSJD-GBM-002 CRISPR cells were cultured in NeuroCul NS-A Basal Medium (Stemcell Technologies, 05750) supplemented with 10% NeuroCult Proliferation Supplement (Human) (Stemcell Technologies, #05753), 20 ng/μL human EGF (Shenandoah, #100-26), 20 ng/μL human FGF-BASIC-154aa (Shenandoah, #100-146), 2 μg/mL Heparin Solution, and penicillin–streptomycin (10,000 U/mL; Thermo Fisher Scientific, #115140122).

### Cell lysate preparation and histone extraction

For whole-cell protein extraction, cells were washed with phosphate-buffered saline (PBS) (Gibco #10010-023) and the pellet formed after centrifugation was resuspended with RIPA lysis buffer (Thermo Fisher Scientific #8990) supplemented with a cocktail of protease (Sigma #P8340) and phosphatase (APExBIO #K1012) inhibitors. After lysis under rotation at 4°C for 30 minutes, the lysate mix was centrifuged, and the supernatant containing whole-cell protein was transferred to a new tube.

For histone extraction, the cell pellet was washed with PBS, resuspended in 1 ml hypotonic lysis buffer (10 mM Tris HCl pH 8.0, 1 mM KCl, 1.5 mM MgCl2 and 1X protease and 1X phosphatase inhibitors), and incubated for 30 min on a rotator at 4 °C. The pellet was collected by centrifugation at 10,000g, 4 °C for 10 min. Then, it was resuspended in 400 μl of 0.4 N H2SO4 and placed on a rotator at 4 °C overnight. Following centrifugation, the supernatant was transferred to a new tube, 132 μl trichloroacetic acid (Sigma #T0699) was added, and the mixture was incubated on ice for 30 minutes. The histone pellet was obtained by centrifugation at 16,000g, 4 °C for 10 minutes and washed with cold acetone twice. Centrifugation at 16,000g at 4 °C for 5 minutes was performed to remove the acetone after each wash. After the last acetone wash, the histones were air-dried at room temperature for 20 minutes before being resuspended in an appropriate volume of ddH2O.

### Immunoblotting

Cells were lysed in 1× RIPA buffer containing 1× protease (100× stock, Sigma Aldrich, P8340) and 1× phosphatase inhibitors (100× stock, Sigma Aldrich, P5726). Protein concentrations were quantified using colorimetric bicinchoninic acid assay (BCA) (Pierce BCA protein assay, 23227). Equal amounts of total protein from cell lysate or histone extracts were loaded on 4–15% Mini-Protean TGX precast gels (Bio-Rad, 3450027). Proteins were transferred to PVDF membranes using a Bio-Rad Trans-Blot Turbo transfer system (Bio-Rad, 1704150). Membranes were blocked with 5% bovine serum albumin (BSA) dissolved in TBST (TBS buffer containing 0.1% Tween-20) and incubated in primary antibody diluted in 5% BSA at 4 °C overnight.

The following antibodies were used in the immunoblotting experiments: H3K27Lac (1428, PTM BIO), H3K9Lac (1419RM, PTM BIO), H3K14Lac (1414RM, PTM BIO), H3K18Lac (1406RM, PTM BIO), H4K12Lac (1411RM, PTM BIO), H4K8Lac (1415RM, PTM BIO), H2BK15Lac (1426RM, PTM BIO), H2BK16Lac (1424RM, PTM BIO), H2AK11Lac (1422RM, PTM BIO), H3K27Cr (545RM, PTM BIO), total H3 (3638, Cell Signaling Technology), total H4 (2592, Cell Signaling Technology), total H2A (12349, Cell Signaling Technology), total H2B (12364, Cell Signaling Technology), H3K27M (74829, Cell Signaling Technology), H3K27me3 (9733, Cell Signaling Technology), H3K27ac (177178, Abcam), CDKN2A/p16INK4a (211542, Abcam), NRG1 (53104, Abcam), Vinculin (V9264, Sigma-Aldrich), NG2 (43916, Cell Signaling Technology), GAPDH (2118, Cell Signaling Technology), OLIG2 (65915, Cell Signaling Technology), GFAP (12389, Cell Signaling Technology), HIF1a (179483, Abcam), beta-actin (4967, Cell Signaling Technology), LDHA (2012, Cell Signaling Technology), LDHB (52488, Abcam), CBP (7389, Cell Signaling Technology), p300 (86377, Cell Signaling Technology), PTEN (9559, Cell Signaling Technology), HK2 (sc-374091, Santa Cruz Bio), ACSS2 (ab133664, Abcam).

Next, membranes were washed with TBST three times. Membranes were incubated with species-matched secondary antibodies (1:5,000), goat-anti-mouse (Bio-Rad, 1706516) or goat anti-rabbit (Bio-Rad, 1706515) conjugated to horseradish peroxidase (HRP) for 2 h at room temperature and washed with TBST 3 times (about 5 min each) and TBS one time (around 10 min). Immunoreactivity was detected using Pierce ECL Western blotting substrate (Thermo Fisher Scientific, 32106)

### Gene silencing

Target gene knockdown was performed using predesigned MISSION shRNA lentiviral plasmids obtained from Sigma-Aldrich. The specific clones used are as follows: shH3-3A (clone ID: TRCN0000139066), shCBP (clone IDs: TRCN0000006485 = sh1 or TRCN0000006486 = sh2), shp300 (clone IDs: TRCN0000009883 = sh1 or TRCN0000039885 = sh2), shACSS2 (clone IDs: TRCN0000045563 = sh1 or TRCN0000045564 = sh2), shLDHA (clone ID: TRCN0000026538), and shLDHB (clone ID: TRCN0000028502). H3.3WT, H3.3K27M, H3.3G34R lentiviral plasmids were obtained from the late David Allis, The Rockefeller University.

shRNA-containing lentiviral particles were generated according to the LentiStarter 3.0 kit (SBI #LV060-1, Palo Alto, CA). Briefly, HEK293T cells were transfected with 700 ng of human shH3-3A plasmid purchased from Sigma (clone ID: TRCN0000139066).

HEK293T-transfected media containing lentiviral particles were collected 72 h post transfection and filtered with a 0.45 µm-syringe filter. Next, the filtered lentiviral particles were added into media of previously plated mNSCs, DIPG007, or R059 cells. After 48 h post transduction, culture media was replaced by fresh growth media. After an additional 48 h, media was supplemented with 1 μg/mL puromycin for selection of successfully transduced cells. Knockdown was confirmed via immunoblotting.

### Chemical Compounds

<u>CBPD-409</u> proteolysis-targeting chimera (PROTAC) compound was provided by Shaomeng Wang (University of Michigan). Cells were treated with 5 nM, 10 nM, or 100 nM CBPD-409 for 48 hr, unless otherwise noted. Dimethyl sulfoxide (DMSO) (Sigma #D2650) was used as vehicle.

<u>NCI-006</u>, lactate dehydrogenase (LDH) inhibitor, was provided by Bryan Mott (Wake Forest University). Cells were treated with 1 mM or 5 mM NCI-006 for 48 hr, unless otherwise noted. DMSO (Sigma #D2650) was used as vehicle.

<u>GSK2837808A</u>, commercially available lactate dehydrogenase A (LDHA) inhibitor, was purchased from MedChemExpress (Cat #HY-100681). Cells were treated with 1 mM, 2 mM, or 5 mM GSK2837808A for 48 hr, unless otherwise noted. DMSO (Sigma #D2650) was used as vehicle.

<u>MTB9655</u>, commercially available acetyl-Coenzyme A Synthetase 2 (ACSS2) inhibitor, was purchased from Selleck Chemicals (Cat #E1211). Cells were treated with 1 uM or 5 uM MTB9655 for 48 hr, unless otherwise noted. DMSO (Sigma #D2650) was used as vehicle.

<u>L-lactic acid</u> was purchased from MedChemExpress (Cat #HY-Y0479). Cells were treated with 2 mM, 5 mM, 10 mM, or 15 mM L-lactic acid for 48 hr, unless otherwise noted. PBS (Gibco #10010-023) was used as vehicle.

### Condition medium

Cells were plated and allowed growth to a confluency of >50% before performing a media change. After an additional 48 hours of growth, the medium was removed and filtered through a 0.45μm polyethersulfone membrane. Then, the medium was boiled at 100°C for 15 minutes before centrifugation to remove any remaining precipitate. Then, the filtered, boiled, and centrifuged conditioned medium was resuspended in the appropriate growth medium to the original volume. Conditioned media was used at a 3:1 ratio with fresh media to avoid effects of nutrient exhaustion.

### Serum differentiation

Freshly dissociated patient-derived H3K27M cells (DIPG007 and BT245) were cultured in complete growth medium supplemented with 10% Fetal Bovine Serum (FBS) (Cat#VWR #89510-186) for 2 weeks to establish serum-differentiated cell lines. During the 2-week timeframe, cells were passaged and re-seeded accordingly before reaching 70% confluency to maintain optimal cell viability. After 2 weeks, the serum-differentiated cells were used for experiments.

### Mouse brain dissociation and primary OPCs and ACs isolation

Pregnant CD-1 mice (Charles River Laboratories) were used to obtain embryonic day 18 (E18) pups. Embryonic brains were collected at E18, and the brain tissues were dissociated using the Brain Tumor Dissociation Kit (P) (Miltenyi Biotec, Cat #130-095-942) and the gentleMACS Octo Dissociator with Heaters. Dissociated cells were then further processed using either the anti-ACSA-2 MicroBead Kit, mouse (Miltenyi Biotec, Cat #130-097-678) for ACs isolation, or the CD140a (PDGFRα) MicroBead Kit, mouse (Miltenyi Biotec, Cat #130-101-547) for OPCs isolation. OPCs and ACs were isolated according to the manufacturer’s protocols.

### Lactate quantification

Lactate levels in suspension cell samples (DIPG007 and BT245) and their corresponding cell culture medium were measured using the L-Lactate Assay Kit (Colorimetric; Abcam, Cat. No. ab65331) according to the manufacturer’s protocol. Cell samples were processed with the deproteinization protocol. Lactate concentrations were determined by colorimetric detection using BioTek Synergy HTX Multi-Mode microplate reader (Agilent company).

### TGFβ1 Elisa

TGF-β1 levels in cell culture media were quantified using the Human/Mouse/Rat/Porcine/Canine TGF-β1 Quantikine ELISA Kit (R&D Systems, Cat. No. DB100C) according to the manufacturer’s protocol. Latent TGF-β1 was activated using Hydrochloric acid (HCl) and HEPES-containing Sodium Hydroxide (NaOH). TGF-β1 levels were determined by colorimetric detection using BioTek Synergy HTX Multi-Mode microplate reader (Agilent company).

### Cell Counting

Proliferation and viability of cells after drug treatment was assessed by trypan blue exclusion assays. Cells were seeded at a density of 2.5 × 10^4^ cells per well in a 24-well plate and were left overnight. They were then treated with vehicle or drug (CBPD409) at varying concentrations and time points for the indicated time periods, following which they were counted using a Countess III cell counter (Thermo Fisher Scientific).

### Subcutaneous H3K27M DMG xenograft model

Flank models of H3K27M DMG were established in NSG mice by injecting 2∗10^6^ patient-derived DMG cells (R059 or UMPED83) in PBS mixed with equal volume of Matrigel in flanks of mice. Treatment started after palpable tumors (around 200mm^3^) appeared. Mice were treated with vehicle or 3mg/kg CBPD-409 by oral gavage for 2 weeks, 3 days per week. Tumor volumes were measured twice a week using Vernier calipers. Tumor volumes were calculated using the formula L (longer dimension) × W (shorter dimension) × W/2. Animals were humanely euthanized at the study’s endpoint before collecting the tumor tissues.

### [13C]-Isotope tracing and analysis

#### In vitro tracing

For ^13^C tracing experiments, 2 million cells were plated and cultured for 16 h in medium supplemented with either ^13^C-U-labelled glucose (25 mM) (2 mM; Cambridge Isotope Laboratories, CLM-99-1396-1). Cells were collected by centrifugation, and the culture medium was aspirated. Cell pellets were resuspended in 300 μl ice-cold methanol by thorough pipetting. Subsequently, 300 μl water containing 1 μg norvaline as an internal standard and 600 μl chloroform were added to each microfuge tube.

Samples were thoroughly vortexed for 30 min at 4 °C and then centrifuged. The polar layer was collected, dried in a SpeedVac for 3–4 h without heat, and stored at −80 °C. For derivatization, 30 μl methoxyamine hydrochloride (Fisher Scientific, PI45950) was added to the dried samples, which were incubated at 45 °C for 30 min with constant mixing. Next, 30 μl MBTSTFA containing 1% TBDMCS (Sigma-Aldrich, M-108-1243, 5 × 1 ml) was added, and samples were incubated at 45 °C for 1 h. Derivatized samples were then transferred to GC vials containing glass inserts.

Samples were analyzed using an Agilent 7890 gas chromatograph equipped with a 30-m HP-5MSUI capillary 1246 column coupled to an Agilent 5977B mass spectrometer operated in scan mode. A 1–2 μl sample was injected at 270 °C using helium as the carrier gas at a flow rate of 1 ml min^−1^. The temperature gradient was maintained at 100 °C for 1 min, increased at 3.5 °C min^−1^ to 255 °C, then increased to 320 °C at 15 °C min^−1^ and held for 3 min, for a total method time of 52.6 min. The MS detector was operated in scan mode over a range of 70–600 *m/z*.

Relative metabolite abundances were calculated by normalizing the integrated peak area of each ion to the norvaline internal standard signal and protein content. For ^13^C-labelled samples, raw GC–MS peak data for all reported metabolites were subjected to isotopic correction using the IsoCorrectoR package (v.1.5.1), available through the Bioconductor library (BioC 3.8), and implemented in R (CRAN 3.6.1).

#### In vivo tracing

At 2-3 weeks following orthotopic tumor implantation, mice bearing intracranial DIPG007 or PPK tumors underwent dual catheterization. One catheter was surgically placed in the jugular vein for tracer administration, and a second catheter was placed in the carotid artery for plasma collection during tracer infusion. Mice were allowed to recover from surgery for 4–5 days. Conscious, freely moving, and undisturbed mice were subsequently administered stable isotope tracer through the intravenous catheter. [U^13^C]-glucose was administered as a bolus dose of 0.4 mg g^−1^, followed by continuous infusion at 12 ng g^−1^ min^−1^ for a total of 4 h.

During the infusion period, blood samples were collected periodically through the arterial catheter into EDTA-coated vials and subsequently processed to obtain plasma. At the completion of the infusion, ketamine (50 mg kg^−1^) was administered through the intravenous catheter to rapidly induce anesthesia. Mice were then decapitated, and tissues were collected on dry ice. To separate orthotopic diffuse midline glioma (DMG) tissue from the mouse cortex, microdissection was performed under fluorescent illumination to distinguish GFP-positive tumor tissue from GFP-negative cortical tissue. All tissues were immediately flash-frozen in liquid nitrogen, within 3 min of anesthesia, for subsequent analysis.

### Magnetic Resonance Spectroscopy (MRS)

MRS was performed as previously outlined (Panigrahy et al., 2006), along with standard pre- and post-contrast MR imaging on 1.5T and 3T MR systems (Signa LX, GE Healthcare, Milwaukee, Wisconsin). All acquisitions employed single-voxel point-resolved spectroscopy (PRESS) with a short echo time (TE) of 35 ms, a repetition time (TR) of 1.5 seconds, and 128 signal averages. The sizes and shapes of the regions of interest (ROIs) were tailored to the size of the lesions, typically ranging from 5 to 10 cm³. The overall acquisition time, including scanner adjustments, was under five minutes per spectrum. Spectra were analyzed using the fully automated LCModel software (Stephen Provencher Inc., Oakville, Ontario, Canada, LCModel Version 6.3– 1L) (Provencher, 1993). T2-weighted fast spin-echo, FLAIR, and T1-weighted FLAIR images were obtained in all cases, with the ROI documented across at least three MR images. The MRS data evaluation focused on metabolites that have been consistently measured and reported in previous research (Bluml et al., 2011; Panigrahy et al., 2006), including lactate (Lac), choline (Cho), myoinositol (mI), and creatine (Cr). Absolute concentrations (institutional units) were calculated using the unsuppressed water signal as a reference and the default water content set by the LCModel software (65%).

### RNA-sequencing

RNA was isolated using Trizol (Invitrogen 15596-026) and treated with DNase (Sigma, 9003-98-9). RNA-sequencing was then performed by the Advanced Genomics Core at the University of Michigan. RNA was assessed using the TapeStation (Agilent) and Qubit RNA broad-range assay (Thermofisher). Samples with RINs (RNA Integrity Numbers) of 8 or greater were prepared using the NEBNext Poly(A) mRNA Magnetic Isolation Module (NEB), xGen Broad-range RNA Library Prep (IDT), and xGen Normalase UDI Primer Plate 2 (IDT), where 200ng of total RNA was converted to mRNA using polyA purification. The mRNA is then fragmented and copied into first strand cDNA using reverse transcriptase and random primers. The 3 prime ends of the cDNA are then adenylated, and adapters are ligated. The products are purified and enriched by PCR to create the final cDNA library. Final libraries were checked for quality and quantity by Qubit hsDNA (Thermofisher) and LabChip (Perkin Elmer). The samples were pooled and sequenced on the Illumina NovaSeq S4 Paired-end 150bp, according to manufacturer’s recommended protocols. Reads were processed and analyzed as described in “Bioinformatics analysis” section of methods shown below.

### ChIP-sequencing

ChIP experiments were performed per the manufacturer’s protocol using an Ideal ChIP– seq kit for histones (Diagenode, Cat#C01010051). For histone ChIPs, 1 million cells were used per ChIP reaction with 2 µg antibody. In brief, cells were crosslinked for 10 min in a 1% formaldehyde solution, followed by quenching with 1/10 the volume of 1.25 M glycine for 5 min at room temperature. Following this, the cells were lysed and sonicated (Bioruptor Pico, Diagenode) to a desired chromatin size of about 200 bp using the Easy mode cycle. Sheared chromatin was then incubated with the following antibodies overnight at 4 °C: H3K27Lac (PTM BIO, Cat# PTM-1428) and H3K27ac (Diagenode, Cat# C15410196). ChIP DNA was de-crosslinked and purified the next day using a Diagenode iPure kit V3 following the manufacturer’s protocol. ChIP-seq libraries were made by using NEB Multiplex dual barcodes (E6448L) and KAPA Hyper Prep kit(96rxn)(Roche Diagnostics KK8504) according to manufacturer’s protocol. Briefly, 6-100ng ChIP DNA samples were end repaired and A-tailed, then ligated with NEB adapters, following by 2xKAPA HiFi HotStart mix and NEB dual indexes PCR to enrich fragments between 200-500bp. Library quantification and quality were measured using Agilent 2100 Bioanalyzer (DNA 1000 chip). Sequenced by Illumina NovaSeq. Run parameters were: 151|8|8|151. Reads were processed and analyzed as described in “Bioinformatics analysis” section of methods shown below.

### Bioinformatics analysis

#### H3K27Lac-dominant versus H3K27ac-dominant

For each cell line and condition, H3K27Lac and H3K27ac MACS3 narrowPeak calls were merged into a unified set using pybedtools. Paired-end fragments from deduplicated BAM files were counted over these regions using featureCounts (-p --countReadPairs -B -O) and normalized to counts per million (CPM). Relative mark enrichment across autosomal peaks (chr1–22) was defined as log2[(CPM_H3K27Lac + 0.01)/(CPM_H3K27ac + 0.01)], with regions scoring >1 or <-1 classified as H3K27Lac- or H3K27ac-dominant, respectively.

### ChIP-seq preprocessing and alignment

Paired-end ChIP-seq FASTQ files were received already adapter-trimmed. Read quality was assessed with FastQC (v0.12.1). Reads were aligned with Bowtie2 (v2.4.2) using --local --very-sensitive-local with a fragment insert-size range of 10–700 bp (-I 10 -X 700). Human samples were aligned to GRCh38, and mouse samples were aligned to mm10. Effective genome sizes used for RPGC normalization were 2,820,131,061 (GRCh38) and 2,419,004,802 (mm10). Bowtie2 output was streamed through SAMtools (v1.21) for BAM conversion, coordinate sorting, and indexing. Duplicates were marked with Picard MarkDuplicates (v3.0.0). Duplicate-marked BAMs were then filtered with samtools view -F 1804 -f 2 -q 10, retaining properly paired reads with mapping quality ≥10 and excluding unmapped reads, mate-unmapped reads, secondary alignments, QC-failed reads, and duplicates. Duplicate-removed BAMs were re-indexed and used for all downstream analyses. Alignment and library quality were summarized with samtools flagstat and samtools stats. Fragment-size distributions were computed from a 10% random subsample of read pairs (samtools view -s 0.1), retaining read pairs with an inferred fragment length (|TLEN|) between 1 and 2,000 bp. RPGC-normalized BigWig files were generated with deepTools (v3.5.6) bamCoverage using --binSize 10 and the species-appropriate effective genome size. These BigWig files were used for genome-browser visualization and as input to deepTools heatmap and profile analyses.

### Peak calling

Narrow peaks were called with MACS3 (v3.0.3) in paired-end mode (-f BAMPE) using pre-pooled IgG control BAMs. IgG samples were pooled per cell line for BT245 (human) and mNSCs (mouse), and per cell-line pair for DIPG007 and SF188 (which shared a single human IgG pool). Effective genome sizes matched those used for RPGC normalization (2,820,131,061 for human, 2,419,004,802 for mouse). Peaks were called at q < 0.05 with --call-summits; --keep-dup all was set because duplicates had already been removed during preprocessing. Peaks were annotated with HOMER annotatePeaks.pl (HOMER v4.11.1) against the matched reference build (hg38 for DIPG007, BT245, and SF188; mm10 for mNSCs), with genomic-feature annotation statistics written via -annStats.

### Genomic region classification of ChIP-seq peaks

Per-sample MACS3 narrow peaks were classified against a cell-line-specific reference of regulatory regions built from H3K27Ac peaks. H3K27Ac served as the reference mark, and H3K27Lac peaks were characterized relative to this H3K27Ac reference. For each cell line, all available H3K27Ac MACS3 narrowPeak files were combined into a single GRanges object and merged using GenomicRanges::reduce with min.gapwidth = 101, such that peaks separated by ≤100 bp were merged. Candidate enhancer regions were defined as merged intervals supported by H3K27Ac peaks in at least two samples for BT245, DIPG007, and mNSCs (each contributing five conditions: H3K27M parental, H3WT-like, lactate, and CBPD), and for one sample in SF188 (single H3WT sample).

Promoter regions were defined as the ±1 kb window around each annotated transcription start site, obtained via promoters(genes(txdb), upstream = 1000, downstream = 1000). The TxDb package was selected per cell line (TxDb.Hsapiens.UCSC.hg38.knownGene (v3.20.0) for human lines, TxDb.Mmusculus.UCSC.mm10.knownGene (v3.10.0) for mNSCs). Peaks overlapping any promoter region were labeled “promoter,” non-promoter peaks overlapping any candidate enhancer regions were labeled “enhancer-like,” and all remaining peaks were labeled “other.” Reference construction and per-sample classification were implemented in R (v4.4.3) via rpy2 (v3.6.4) using Bioconductor (v3.20) packages GenomicRanges (v1.58.0), rtracklayer (v1.66.0), and GenomeInfoDb (v1.42.3). Analyses were run independently for each cell line.

### Quantification of ChIP-seq peaks

#### Counts per million

For each cell line (DIPG007, BT245, mNSCs, SF188), a union peak set was built by pooling MACS3 narrowPeak calls from all conditions and both H3K27Lac and H3K27Ac marks, restricting to canonical autosomes (chr1–22 for hg38; chr1–19 for mm10), sorting, and merging overlapping intervals with bedtools (v2.31.1, via pybedtools). Each peak was annotated with boolean columns indicating the condition × mark combinations in which the peak was called, so pairwise comparisons could be restricted to peaks called in the conditions being compared. Read pairs were counted over the union peak set with Subread featureCounts (v2.0.3; Liao et al., Bioinformatics 2014) in paired-end fragment mode (-p --countReadPairs -B); the -O multi-overlap flag was disabled because the union is non-overlapping by construction. All H3K27Lac and H3K27Ac duplicate-removed BAMs for a cell line were quantified in a single invocation, yielding one peaks by sample count matrix. Library sizes were defined as mapped primary read pairs in the duplicate-removed BAM (samtools view -c -f 64 -F 3852; samtools v1.21).

Raw counts were converted to CPM as (count / library size) × 10[and log-transformed as log₂(CPM + 1). Peaks from the union peak set were annotated for nearest gene and genomic feature class with HOMER annotatePeaks.pl (HOMER v4.11.1; Heinz et al., Mol Cell 2010) against the appropriate reference genome and annotations were merged onto the count and CPM tables by peak identifier.

#### Peak signal

H3K27Lac and H3K27Ac peaks were paired at overlapping intervals. For each cell line and condition, MACS3 narrowPeak files were restricted to canonical autosomes and intersected with bedtools intersect -wa -wb (v2.31.1, via pybedtools), using H3K27Lac as the anchor (-a) and H3K27Ac as the overlay (-b). Peaks with multiple overlap partners formed one row per pairing. Per-condition tables were concatenated across BT245, DIPG007, and mNSCs. Each paired peak was labeled by regulatory class (promoter, enhancer-like, or other) as described above.

For the H3K27M parental Lac-versus-Ac scatter plot, pairs were filtered to q < 0.05 at both marks, and raw signalValue was used as the comparison metric. For each cell line, signed residuals from y = x were computed as r = signalValue_Lac − signalValue_Ac, and σ was taken as the sample standard deviation of r. Peaks were classified as H3K27Lac-dominant (r > +0.5 σ), H3K27Ac-dominant (r < −0.5 σ), or co-dominant otherwise. The y = x line serves as a reference axis for visualizing dominance, not as a fitted relationship.

### Analysis of repeat elements

#### Genome-wide H3K27Ac and H3K27Lac deposition

Genome-wide deposition was quantified using a tiling-based strategy adapted from Krug et al. (2019). For each ChIP-seq sample, a HOMER tag directory was generated from the duplicate-removed BAM using makeTagDirectory (HOMER v4.11), and genome-wide tag distributions were computed with annotatePeaks.pl using none as the peak argument, -d <tagDir>, and -annStats. In this mode, HOMER reports per-bin tag counts across its default genome-wide tiling (∼1 Mb windows on hg38 and mm10) together with feature-level summary statistics. Per-window tag counts were converted to CPM using total input read pairs from the corresponding Bowtie2 alignment log as the denominator. For each human cell line (DIPG007, BT245), histone mark (H3K27Ac, H3K27Lac), and pairwise comparison (H3K27M parental versus matched (1) shH3-3A/CRISPR-corrected H3WT, (2) added exogenous lactate, or (3) CBPD treatment), per-window CPM distributions were compared by two-sided Mann–Whitney U test and visualized as boxplots.

#### H3K27Ac and H3K27Lac at repeat element classes

ChIP signal at annotated repetitive elements was quantified from the same annotatePeaks.pl output, using per-element entries from HOMER’s RepeatMasker-derived annotation of the corresponding reference genome (hg38 or mm10). HOMER-normalized tag counts were used as the per-element ChIP signal, and the broad repeat class of each element was parsed from the PeakID field. Following Krug et al., analysis was restricted to the four major classes: LINE, SINE, LTR, and DNA. Within each combination of cell line, condition, and histone mark, per-element signal distributions for each repeat class were visualized as boxenplots, with H3K27M from the matched cell line plotted alongside. Differential signal between H3K27M and control was assessed by two-sided Mann–Whitney U test.

### Heatmap and profile visualization

For multi-mark profiling in H3K27M mNSCs, six histone marks (H3K27Lac, H3K27Ac, H3K27me3, H3K4me1, H3K4me3, H3K36me3) were profiled over the union of their MACS3 narrowPeak calls (n = 287,487 intervals). computeMatrix was run once across all six RPGC-normalized BigWig files in reference-point mode with ±3 kb flanking, and the matrix was rendered with a custom Python script. Rows were restricted to canonical mouse autosomes plus and sorted in descending order by Gaussian-weighted center signal of H3K27Lac, and the same row order was applied to all six columns. Each mark was plotted with its own sequential colormap.

For pairwise condition comparisons within a single histone mark in the human cell lines, two-column matrices (H3K27M parental | condition) were generated over the union of MACS3 narrowPeak calls from the two conditions. Using the peak-level CPM values from Quantification of ChIP-seq peaks section above, peaks were classified by log2 CPM fold change, and grouped as condition-high (log₂FC > +0.5σ), H3K27M-high (log₂FC < −0.5σ), or similar (|log₂FC| ≤ 0.5σ), where σ is the standard deviation of log₂FC across peaks. These results are visualized as heatmaps showing the condition-high (red) and H3K27M-high (blue) heatmaps. Rows within each heatmap were sorted as above. Similar peaks are not displayed but are retained in the denominator for per-group percentages.

### RNA-seq

DIPG007 cells were profiled by bulk paired-end RNA sequencing in three biological replicates across five conditions: parental H3K27M, shH3-3A, 5 mM exogenous lactate (lactate), and 100µM CBPD; SF188 was included as a H3WT pediatric glioma control. FASTQ files were processed at the University of Michigan Advanced Genomics Core with nf-core/rnaseq v3.19.0 on Nextflow v24.10.5. Read quality was assessed with FastQC and MultiQC, adapters were trimmed with Trim Galore (cutadapt), and reads were aligned to GRCh38 with STAR v2.7.10a. Ensembl release 113 gene annotation was used. Gene- and transcript-level expression was quantified with RSEM v1.3.1, the gene count matrix was annotated via BioMart, and alignment and library QC metrics were generated with RSeQC, Qualimap, Picard, and samtools.

#### Differential expression

Pairwise differential expression was performed against DIPG007 parental as the reference in five comparisons: lactate, CBPD, shH3-3A, and SF188. RSEM transcript files were imported with tximport (v1.34.0) and summarized to gene level using a tx2gene table built from the RSEM transcript-to-gene mapping. Gene counts were analyzed with DESeq2 (v1.46.0).

#### Integrated ChIP–RNA gene sets

For each perturbation (lactate, CBPD), integrated ChIP–RNA gene sets were defined as the top 500 genes that changed concordantly in both data modalities versus parental.

Concordance was defined as same-sign log2 fold change in H3K27Lac ChIP-seq and RNA-seq (RNA-seq DESeq2 padj < 0.05). Concordant genes were then ranked by H3K27Lac ChIP-seq log2 fold change and the top 500 retained. A pool size of 500 was used to provide a sufficiently large gene set given the relatively noisier ChIP-seq signal. We considered the gene sets defined by lactate UP/UP, CBPD UP/UP, and CBPD DOWN/DOWN for further analyses.

### Spatial transcriptomics

#### Datasets

Visium spatial gene expression data were obtained from Collot et al.^28^: three human H3K27M DMG samples (VUMC_11_pons, VUMC_17_pons, BT01_192_core) and four mouse H3K27M cortex tumors generated by in utero electroporation s(IUE_CORTEX_TUMOR_B_C1, B_D1, C_A1, C_B1). Each sample was processed from its Space Ranger outs/ directory using the filtered feature-barcode matrix, the high-resolution H&E image, and the corresponding spot scale factors.

#### Preprocessing

Each sample was processed independently in Python with Scanpy (v1.10) and Squidpy. Filtered count matrices were loaded with squidpy.read.visium, gene names were de-duplicated, mitochondrial content was annotated with the species-appropriate prefix (MT- for human, mt- for mouse), and QC metrics were computed with sc.pp.calculate_qc_metrics. Spots with fewer than 200 detected genes and genes detected in fewer than 10 spots were dropped. Counts were library-size normalized to 10,000 transcripts per spot and log1p-transformed.

#### Gene-set scoring

Per-spot gene-set scores were computed on the log-normalized matrix with sc.tl.score_genes, which compares mean expression of the query set against an expression-matched random reference. Two gene-set families were scored on each sample: (1) the MSigDB Hallmark HYPOXIA set, retrieved with gseapy.get_library(“MSigDB_Hallmark_2020”) and (2) the integrated ChIP–RNA gene sets defined above (lactate UP/UP, CBPD UP/UP, CBPD DOWN/DOWN). For mouse samples, gene sets defined on human symbols were converted to mouse orthologs prior to scoring using the Mouse Genome Informatics human–mouse homology report (HOM_MouseHumanSequence.rpt). Genes without a one-to-one ortholog or absent from the sample’s expression matrix were dropped.

#### Spatial correlation analysis

Gene signatures were defined from genes showing concordant changes in H3K27Lac ChIP-seq and RNA-seq (RNA-seq adjusted P < 0.05). For each signature, genes were ranked by the geometric mean of their ChIP-seq and RNA-seq absolute log2 fold-change percentile ranks, and the top 500 genes were retained. Spot-level signature scores were computed on normalized, log-transformed data using Scanpy’s tl.score_genes. Correlations between signature scores and cell-state or hypoxia gene sets were evaluated across quality-filtered spots using Spearman’s rank correlation, calculated independently for each human (n = 3) and mouse (n = 4) sample.

### Pathway analysis

Gene Ontology Biological Process over-representation was tested with the MSigDB “Investigate Gene Sets — Compute Overlaps” tool (https://www.gsea-msigdb.org/gsea/msigdb/human/annotate.jsp), a hypergeometric overlap test with Benjamini–Hochberg false discovery rate (FDR) correction across the queried collection. Queries were run against the C5: GO:BP subcollection and thresholded at FDR < 0.05.

